# A robust evaluation of TDP-43, poly GP, cellular pathology and behavior in a AAV-C9ORF72 (G_4_C_2_)_66_ mouse model

**DOI:** 10.1101/2024.08.27.607409

**Authors:** Emily G. Thompson, Olivia Spead, S. Can Akerman, Carrie Curcio, Benjamin L. Zaepfel, Erica R. Kent, Thomas Philips, Balaji G. Vijayakumar, Anna Zacco, Weibo Zhou, Guhan Nagappan, Jeffrey D. Rothstein

## Abstract

The G_4_C_2_ hexanucleotide repeat expansion in *C9ORF72* is the major genetic cause of both amyotrophic lateral sclerosis (ALS) and frontotemporal dementia (FTD) (C9-ALS/FTD). Despite considerable efforts, the development of mouse models of C9-ALS/FTD useful for therapeutic development has proven challenging due to the intricate interplay of genetic and molecular factors underlying this neurodegenerative disorder, in addition to species differences. This study presents a robust investigation of the cellular pathophysiology and behavioral outcomes in a previously described AAV mouse model of C9-ALS expressing 66 G_4_C_2_ hexanucleotide repeats. Despite displaying key molecular ALS pathological markers including RNA foci, dipeptide repeat (DPR) protein aggregation, p62 positive stress granule formation as well as mild gliosis, the AAV-(G_4_C_2_)_66_ mouse model in this study exhibits negligible neuronal loss, no motor deficits, and functionally unimpaired TAR DNA-binding protein-43 (TDP-43). While our findings indicate and support that this is a robust and pharmacologically tractable model for investigating the molecular mechanisms and cellular consequences of (G_4_C_2_) repeat driven DPR pathology, it is not suitable for investigating the development of disease associated neurodegeneration, TDP-43 dysfunction, gliosis, and motor performance. Our findings underscore the complexity of ALS pathogenesis involving genetic mutations and protein dysregulation and highlight the need for more comprehensive model systems that reliably replicate the multifaceted cellular and behavioral aspects of C9-ALS.

## Introduction

Amyotrophic lateral sclerosis (ALS) is a progressive and ultimately fatal neurodegenerative disease that affects upper and lower motor neurons in the cerebral cortex, brainstem and spinal cord. The degeneration of motor neurons leads to muscle denervation and atrophy with paralysis and death three to five years after disease onset due to respiratory insufficiency [41, 81]. While most ALS cases have no known genetic cause, an intronic GGGGCC hexanucleotide repeat expansion (HRE) in the *C9ORF72* gene is responsible for the majority of familial ALS (fALS) and a subset of sporadic ALS (sALS) [17, 67]. In a healthy individual, the intronic G_4_C_2_ repeat length can range from 2-30 repeats. However, in patients with C9-ALS, this region may contain hundreds to thousands of repeats [7, 17, 31, 47, 67, 71].

The mechanism by which this G_4_C_2_ expansion leads to cellular dysfunction is not yet fully elucidated, however there are several hypotheses currently under investigation. Patients with C9-ALS may show decreased coding C9ORF72 mRNA and protein expression, resulting in loss of function of the endogenous C9orf72 protein [16, 21, 29, 84]. In addition, the repeat expansion causes a gain of toxic function in which the intronic repeat mRNA is transcribed and forms nuclear and cytoplasmic mRNA foci [16, 18, 23]. These cytoplasmic repeat mRNAs can also undergo repeat-associated non-ATG-dependent (RAN) translation, forming aggregation-prone dipeptide repeat proteins (DPRs) [3, 38, 51, 56, 57]. Five different DPR protein species are produced from either sense or antisense transcripts of the HRE. Poly Glycine-Alanine (polyGA), Glycine-Arginine (polyGR), and Glycine-Proline (polyGP) are produced from sense HRE transcripts, while polyGP, Proline-Alanine (polyPA), and Proline-Arginine (polyPR) are generated from antisense HRE transcripts. It is still unclear how or whether RNA foci themselves significantly contribute to cellular toxicity, but it is possible that the RNA foci structures sequester various RNA-binding proteins, leading to the loss of their function [12, 39] and the repeat RNA themselves may impair nucleocytoplasmic transport or the nuclear pore complex structure [14]. DPRs, particularly arginine-rich DPRs, have been implicated in several toxicity-inducing pathways including DNA damage and oxidative stress [44], ER Stress [88], impairment of protein translation and stress granule dynamics [27, 85], and disruption of nucleocytoplasmic transport [20, 32, 73]. Interestingly, in ALS patient brains, DPR accumulations are found in both degenerating and non-degenerating cell types, indicating a more complicated role for DPR inclusions in neuronal toxicity [82].

TAR DNA-binding protein-43 (TDP-43) is a nuclear DNA/RNA-binding protein involved in a several cellular functions including transcriptional as well as post-transcriptional regulation such as RNA splicing, mRNA transport, and translational repression [76]. An overwhelming majority of ALS cases display TDP-43 disruption, characterized by both the nuclear loss of the endogenous protein as well as cytoplasmic inclusions of hyperphosphorylated-TDP-43 [2, 58, 74]. TDP-43 dysfunction due to loss-of-nuclear function and gain-of-cytoplasmic functions are believed to be responsible for TDP-43 driven neurodegeneration [6, 36, 46, 49, 76]. Additionally, TDP-43 aggregates are known to alter many cellular functions through the sequestration of other proteins. For example, in artificial overexpression systems, TDP-43 aggregates sequester nuclear pore complexes and nucleocytoplasmic transport components, altering nuclear import and export [11]. TDP-43 aggregates co-localize with stress granule components, altering their dynamics and possibly increasing cell sensitivity to prolonged stressors [34, 48]. These aggregates are also known to inhibit proteasome function and autophagy which promotes the accumulation of toxic misfolded protein in the cell [22, 53, 58, 77]. It should be noted that recent studies suggest artificial TDP-43 overexpression does not mimic the known changes of TPP-43 loss of function, typically seen in actual human tissue [8].TDP-43 also plays a role in the intracellular and axonal transport of mRNA granules and this function is lost when it is aggregated [1, 19, 72]. However, the actual abundance of TDP-43 aggregate is in reality quite low in human autopsy tissue, while nuclear loss if far more common, suggesting that the loss of function may be the dominant toxicity pathway. While the underlying pathogenic mechanisms mediated by TDP-43 are diverse, it is clear that TDP-43 mislocalization and aggregation lead to neuronal death in C9-ALS [4, 5, 28, 78].

Historically, the use of mouse models has driven *in vivo* efforts to understand disease etiology, mechanisms, and therapeutic opportunities. As with various other neurodegenerative diseases, there has been a concerted effort to create a reliable mouse model for C9-ALS. This has resulted in a plethora of individual models, each with advantages and shortcomings. Collectively, these models have improved our understanding of aspects of C9-ALS pathobiology, such as the role of individual DPRs in pathophysiology. For example, ubiquitous expression of a 50-repeat length polyGR peptide (GR50) resulted in sex-specific neuronal loss and behavioral deficits [83]. Hao and colleagues generated a loxP-Cre line to drive expression of a 28-repeat length polyPR peptide (PR28) in Thy1-neurons, resulting in gliosis, synaptic dysfunction, and motor abnormalities [26]. Similarly, when a 50-repeat length polyGA peptide (GA50) was expressed in the mouse brain using an adeno-associated virus (AAV), mice display neuronal loss throughout the brain and deficits in behavior including memory and motor coordination [89]. While the above-mentioned models sought to express specific DPRs in an attempt to understand specific DPR polypeptide overexpression biology, several models have also been developed to express the G_4_C_2_ HRE at varying repeat lengths, perhaps more closely mimicking authentic human disease. One such model expresses 36-G_4_C_2_ repeats under inducible control with varying phenotypes depending on the promoter [69]. Several groups have used a bacterial artificial chromosome (BAC) DNA clone containing varying lengths of the human *C9ORF72* gene, including G_4_C_2_ repeats ranging from 100-1000, to generate transgenic mice in hopes of modeling “authentic” ALS [30, 42, 60, 61]. Varying degrees of cellular pathology, neuronal loss, and motor phenotypes have been observed in these mouse models. The high variability of expression levels both within and between lines has raised some questions as to the validity of this model. Oftentimes, even replicated studies have had inconsistent results [55, 59].

In addition to germline transgenic models, a mouse model that utilized viral transduction of 66 G_4_C_2_ repeats ((G_4_C_2_)_66_) directly into the CNS of neonatal mice was developed [10, 69]. Upon transduction, neurons express the (G_4_C_2_)_66_ RNA which is translated to produce the three sense-strand-specific DPRs (polyGA, GP, and GR). These mice were found to display many of the pathologies seen in ALS/FTD patients including RNA foci, DPR pathology, pTDP-43 inclusions, and motor deficits as tested by rotarod [10]. It is not yet known whether these exhibit nuclear clearance or loss of function of TDP-43, both typical in authentic human C9-ALS. Moreover, a robust characterization of the AAV-(G_4_C_2_)_66_ mouse model, including comprehensive assessment correlating the biochemical pathology, cellular dysfunction, and behavioral outcomes associated with this model was needed. Therefore, in this study we sought to make this assessment using a robust study design including randomization of animals at multiple levels, blinding until completion of statistical analysis, and a sample size based on power analysis with two primary end points (rotarod and brain polyGP).

## Methods

### Animals

C57BL/6J were purchased from Jackson Laboratories (Strain #000664) at 4-8 weeks of age. At 6-9 weeks of age breeding pairs were established to produce pups for all subsequent experiments. Both male and female mice were used throughout the study and randomly allocated to each of the experimental groups. Mice were housed in a constant 14-hour light/ 10-hour dark cycle and allowed access to food and water *ad libitum*. All animal procedures were carried out in compliance with animal protocols approved by the Animal Use Committee at the Johns Hopkins University School of Medicine (JHUSOM) as well as in accordance with the GSK Policy on the Care, Welfare and Treatment of Laboratory Animals. This study was reviewed by the Institutional Animal Care and Use Committee at GSK, and at JHUSOM where the work was performed.

### Generation of AAV2/9-(G_4_C_2_)_2_ and AAV2/9-(G_4_C_2_)_66_ vectors

The AAV2/9-(G_4_C_2_)_2_ and AAV2/9-(G_4_C_2_)_66_ viruses were provided by Dr. Leonard Petrucelli at Mayo Clinic Jacksonville. The viruses were prepared as previously described [10, 75]. Following generation, the genomic titer of each virus was determined by qPCR and the AAVs were diluted with sterile phosphate buffered saline (PBS) and stored at -80°C until use.

### Randomization, Blinding, and Power Analysis

This study was powered based on rotarod performance and brain polyGP levels as the primary endpoints, motivated by a review of the raw data from the previously described study [10]. The pooled standard deviation of latency-to-fall was estimated as 24.87 seconds with 20 degrees of freedom. In order to obtain a conservative power estimate and mitigate the risk of underpowering the current study, the upper 80% confidence limit of 29.13 seconds was used to calculate the statistical power. The power analysis was based on an ANOVA model accounting for sex and experimental group as factors. With a target sample size of n=20 per experimental group (n=10 per sex per group), this study had 80% statistical power to find statistically significant differences in rotarod response between groups as small as 23.3 seconds.

Similarly, an analogous power analysis was conducted using polyGP data [24], which yielded a standard deviation estimate of 1.66 log-2 cells/µm^2^ with 8 degrees of freedom. Using the upper 80% confidence limit (2.2 log-2 units) of this estimate, this study had 80% statistical power to find statistically significant differences in quantification of polyGP as small as 1.76 log-2 units, or a 3.39-fold increase between groups in cells/µm^2^.

Within the first 24 hours of life (postnatal day 0-1), pups were randomized to receive either AAV2/9-(G_4_C_2_)_66_ injection, AAV2/9-(G_4_C_2_)_2_ injection, or no injection, littermate matched at a ratio of 1-1-1, respectively. Upon weaning, animals were separated by sex and randomized to cages. Animals with the same AAV-(G_4_C_2_)_x_ repeat injection were housed in the same cage. Upon in-life study completion, ex-vivo study samples were randomized among plates for the MSD assay.

Investigators who were monitoring the behavior performance and in-life care were blinded to the experimental groups. Body weight collection throughout the study and end-of-life tissue collection were not blinded. Ex-vivo samples were labeled with only mouse IDs without experimental group information.

### Neonatal Viral Injections

AAV viral aliquots were thawed on ice and spun down in a centrifuge at 4°C. In a sterile hood, viruses were diluted to 1.5x10^10^ viral genomes/µL (vg/µL) with sterile PBS and were stored on ice until time of injection. Intracerebroventricular (ICV) injections of AAV were performed on C57BL/6J postnatal day 0 (P0) pups. AAV dilutions were prepared on the day of injections. Pups underwent cryoanesthesia on ice for approximately 3 minutes or until pups exhibited no movement. A 32-gauge needle (Hamilton; Small RN 32 gauge, 0.5 inch needle, point style 4) attached to a 10 µL syringe (Hamilton, Model 701 RN) was inserted approximately two fifths of the distance between the lambda and each eye at a 30° angle from the surface of the head and was held at a depth of about 2 mm. 2 µL of virus was manually injected into each cerebral ventricle and the needle was held in place for an additional 5 seconds after each injection to prevent back flow. After injections, pups were placed on a heating pad until fully recovered and then returned to their home cages with the dam. Any pups with back flow from the injection were excluded from the study.

### Intracerebroventricular (ICV) anti-sense oligonucleotide (ASO) injection

#### ASO Preparation

A C9ORF72-ASO was prepared by Integrated DNA Technologies with the following sequence: 5’-mC*mC*mG* mG*mC*C* C*C*G* G*C*C* C*C*G* mG*mC*mC* mC*mC -3’ with standard desalting, Na^+^ Salt Exchange, and analytical RP-HPLC. Additionally, to reduce adverse *in vivo* effects from Ca^2+^ chelation, we pre-saturated ASO calcium binding sites by transferring the ASO to a 3 kDa-cutoff ultrafiltration cartridge (Amicon) and washed twice with 10 mM CaCl_2_ followed by water, and finally with PBS [54]. All spins were performed at 4°C. The ASO was then resuspended with PBS, concentration determined with NanoDrop, and diluted to 50 ng/μL with PBS for ICV injection.

#### ICV injection

Delivery of the above described C9-ASO was performed in mice at 90 dpi (days post AAV injection) with stereotactic ICV injection. After isoflurane anaesthesia, slow-release Buprenorphine was injected subcutaneously (1 mg/kg) prior to the surgery. While maintaining anaesthesia via nose cone, the animal’s skull was immobilized on a Stoelting stereotaxic frame. Hair was removed from the scalp with a chemical depilatory cream (Nair) and sterilized with topical disinfectant (Betadine) and 70% ethanol. Using sterile instruments and gloves, a mid-sagittal longitudinal incision of approximately 0.5 cm was made on the scalp to expose the skull and the subcutaneous tissue and periosteum were scraped from the skull with a sterile cotton tipped applicator. At the following coordinates: 0.0 mm anterior to bregma and 1.0 mm to the right lateral, a small burr hole was drilled through the skull with a hand drill attached to a micromanipulator to expose the brain. A Hamilton syringe (Hamilton 7635-01) was filled with either PBS or C9-ASO at 50 ng/μL and attached to a microinjection pump (World Precision Instruments Micro 4) on a stereotactic arm. After alignment of the syringe needle at 90° to the burr hole and surface of the brain, the syringe was slowly lowered 1.9 mm into the brain. 10μL of the loaded solution was then injected with the microinjection pump at a rate of 2 μL/min, followed by a 5-minute wait period. The syringe was then slowly removed, and the incision site was closed with 5-0 polyglactin 910 (Vicryl) absorbable suture. Antibiotic ointment (Fisher Scientific 19-090-845) was applied to the wound and the animal was allowed to fully recover from the anaesthesia on a warmed heating blanket. Carprofen (Rimadyl) at 0.05 mg/mL was administrated in drinking water for seven days post-surgery to manage pain.

### Body Weight measurements

Body weight (g) was measured every two weeks for all animals starting at 90 dpi until the conclusion of the study.

### Behavioral Assays

Mice underwent several behavior tests, including rotarod, grip strength, catwalk, and open-field assessments, at 90 and/or 180 dpi. One week before behavioral assessments, mice were habituated to the equipment used for behavior testing as well as to those running the behavior tests. Animals were placed on the equipment and allowed to explore for two minutes each. Scientists running the tests also gently handled the mice for 2-3 minutes, placing their hand in the cage and allowing the mice to explore, to decrease the stress in the mice prior to behavioral testing.

#### Rotarod

Motor coordination and learning was assessed using an automated rotarod system (Columbus Instruments, Rotamex-5, 0254-8000). Mice were gently placed on a rod that accelerated from 4 to 40 rpm at a rate of 1.2 rpm every ten seconds. Four trials were run for each mouse with at least 20 minutes of rest in between. The latency to fall time was recorded for each trial when the mouse fell from the rod, triggering the sensor to automatically stop the timer. When the mouse fell and clung to the rod or rolled around the rod three consecutive times, the mouse was manually removed from the rod and the corresponding time was recorded as the time of the fall.

#### Grip Strength

The strength of the fore- and hindlimbs of each mouse was measured using a T-bar with an attached meter (Chatillon 2 LBF Ametek Force Gauge) to measure the peak force exerted from each mouse. Each mouse was held by the tail and lowered gently until the front paws were gripping overhand on the T-bar. The mouse was gently pulled back and the force exerted on the bar just before the mouse lost its grip is recorded. For hindlimb assessment, mice were held by the tail and gripped a metal rod with their forelimbs. The mice were lowered until their back paws gripped the T-bar. They were gently pulled back and the maximum force before losing grip was recorded. Each mouse was tested four times for both fore- and hindlimbs.

#### Catwalk

Gait analysis was assessed using the Catwalk XT Automated Gait Analysis System (Noldus Technology, Software Version 10.6.608). Mice walked within a corridor across an illuminated glass platform while a camera recorded underneath. Walking from one end to the other was recorded as one trial and each mouse ran five consecutive trials. From the recordings, each individual pawprint was digitized and labeled into a discrete number of pixels with a range of brightness. This information was quantified using CatWalk (ver. 10.6.608) software to generate numerous parameters of quantitative and qualitative assessment of individual pawprints and gait.

#### Open field

The SDI (San Diego Instruments, San Diego, CA, USA) open field apparatus consisted of a plexiglass square box, 40.6 cm (W) x 40.6 cm (D) x 38.1 cm (L), with side mounted photobeams at 7.6 cm above the floor to measure rearing. All surfaces within the box were thoroughly cleaned with Vimoba disinfectant prior to testing each animal. Mice were placed in the center of the box and allowed to explore the area for 15 minutes. All movements were recorded and analyzed with the Photobeam Activity Systems Data Reporter software (SDI), and the following parameters were measured: total distance traveled, average speed, mobile time, and distance traveled in center zone (20 cm x 20 cm) vs. total distance traveled.

### Statistical Analysis: Behavioral Assays

Rotarod data was analyzed using a two-way ANOVA with timepoint (90 dpi, 180 dpi) and experimental group as factors, controlling for repeated measures on mice. Grip strength at 90 dpi and open-field locomotion activity at 180 dpi were each analyzed using a one-way ANOVA with experimental group as a factor. Body weight was analyzed separately for males and females with a two-way repeated-measures ANOVA with timepoint and experimental group as factors. All ANOVAs were performed in Graphpad Prism (ver. 9.4.1). Overall family-wise error rate was controlled in all post-hoc pairwise comparisons for all assays using the Tukey-Kramer multiple comparison test [37, 79] unless otherwise noted.

The catwalk data at 90 dpi and 180 dpi was analyzed separately at each timepoint, using a linear mixed-effects model to account for correlation among 12 parameters (base of support, phase dispersion, stride length, body speed, cadence, number of steps, run speed, swing speed, print area, stand time, step cycle, and swing time). This no-intercept model used the interaction of experimental group and parameter as fixed effects, in order to obtain the group effect for each parameter; it also used a random intercept for each animal and parameter-specific residual variance estimates. Post-hoc least-squares means were compared among groups within parameter to assess differences in experimental groups. Multiplicity among comparisons for all parameter readouts was controlled using the Tukey-Kramer multiple comparison test. No statistically significant differences were found between any two experimental groups in any of the parameters; three parameters (step number, cadence, run speed) are displayed in **Figure 7D**. The catwalk analysis was performed using SAS (ver. 9.4).

### Cerebrospinal Fluid (CSF), Plasma, and Tissue collection

#### CSF

The mouse was anesthetized with an I.P. injection of 100 mg/kg ketamine-2.5 mg/kg acepromazine-10 mg/kg xylazine. When the animal was no longer responsive to toe pinch the neck musculature was retracted to expose the cisterna magna. A cauterizing iron was utilized to stop any bleeding from the musculature and a cotton applicator soaked in PBS was used to remove any blood. A 25-gauge needle was then used to puncture the cisterna magna dura and a 20 μL pipette was used to collect the CSF (approximately 5-20 μL). Any blood contaminated CSF samples were centrifuged at 3000 x *g* and the supernatant was transferred to a new tube. All samples were stored in low-binding microcentrifuge tubes (Eppendorf 0030108442) on dry ice prior to storage at -80°C.

#### Plasma

After CSF collection, the chest cavity of the animal was opened to provide access to the heart. A 20-gauge needle was attached to a 3 mL syringe and used to puncture the left ventricle of the heart. Negative pressure was slowly applied via the syringe for blood collection. The blood was centrifuged at 3000 x *g* at 4°C for 15 minutes. The plasma supernatant was aliquoted at 60 μL per microcentrifuge tube and snap frozen. All samples were stored at -80°C.

#### Brain

The anesthetized mouse was transcardially perfused with ice-cold PBS containing 10 U/mL heparin (Sigma-Aldrich H3149) for approximately 5 minutes. Subsequently, the brain was removed, and the right hemisphere was snap frozen in a microcentrifuge tube. The left hemisphere was either snap frozen or put into a microcentrifuge tube containing 4% paraformaldehyde in PBS overnight at 4°C and then moved into PBS containing 0.1% sodium azide for long store storage.

#### Spinal Cord

The spinal cord was extracted by hydraulic extrusion, as previously described [68]. The spinal column was dissected, cutting distally to the pelvic bone and proximally of the shoulders, and then trimmed until the spinal cord was visible at both ends. A 20-gauge needle connected to a 5 mL syringe filled with PBS was inserted into the distal end of the spinal column, ensuring the needle was stabilized within the column. Making sure the spinal column was straightened as much as possible, steady pressure was applied to extrude the spinal cord into a petri dish filled with cold PBS. The spinal cord was cut into Cervical, Thoracic, and Lumbar sections. The Thoracic sections were snap frozen and the Cervical and Lumbar sections were put into microcentrifuge tubes containing 4% paraformaldehyde in PBS overnight at 4°C and then moved into PBS containing 0.1% sodium azide for long store storage.

### Brain and Spinal Cord Homogenization

Frozen whole brain hemispheres and thoracic spinal cord samples were quickly transferred to a Lysing Matrix D tube (MP Biomedicals 116913050) and ice-cold Homogenization Buffer (Tris-EDTA buffer, pH 8.0 [ThermoFisher Scientific AM9858] and 1X Protease [Millipore 539131] and Phosphatase Inhibitor Cocktail [Bimake B15002]) was added. The sample was homogenized with an MPbio FastPrep machine for 1 minute. After homogenization the tube was spun at 1500 x *g* for 2 min. at 4°C to remove bubbles. Part of the homogenate was transferred to a tube containing Trizol (ThermoFisher Scientific 15596026) for subsequent RNA isolation, while the rest of the homogenate was transferred to a tube containing 2X Lysis Buffer (50 mM Tris pH 7.5 [Quality Biological 351-006-131], 250 mM NaCl [Quality Biological 351-036-101], 2% Triton X-100 [Sigma-Aldrich T9284], 4% SDS [ThermoFisher Scientific AM9820], 2X Protease [Millipore 539131] and Phosphatase Inhibitor Cocktail [Bimake B15002], and 2 mM PMSF [ThermoFisher Scientific 36978]) and vortexed briefly to mix. The lysate was incubated on ice for 10 min., after which the lysate was spun at 16,000 x *g* for 20 min. at 4°C. The supernatant was transferred to a fresh tube and stored at -80°C until further analysis.

### Meso Scale Discovery (MSD) Assay

#### Sample Preparation

Brain and spinal cord lysate samples were thawed on ice and a BCA Protein Assay (ThermoFisher Scientific 23225) was performed per manufacturer’s instructions to determine total protein concentration of the sample. Subsequently, samples were prepared at 1.0 μg/35μL homogenate buffer and stored at -80°C.

#### MSD Assay

Blocking buffer (Meso Scale Diagnostics R93AA-1) was prepared per manufacturer’s instructions, added to the wells of an MSD Gold 96 Well Small Spot plate (Meso Scale Diagnostics L45SA1), and incubated overnight at 4°C. The plate was washed with PBST (0.05% Triton X-100 in PBS). A polyGP capture antibody (generated in [9]) was conjugated with biotin in house (ThermoFisher Scientific A39256), diluted to 0.375 μg/mL PBS, added to each well, and incubated with shaking at room temperature for 1 hour. The plate was then washed with PBST and poly-(GP)_8_ peptide (CKK-PEG2-GPGPGPGPGPGPGPGP-amide; 21^st^ Century Biochemicals) standards and brain hemisphere or spinal cord samples (1.0 μg/well), diluted in PBS, were added to the plate in triplicates. Incubation occurred for 3 hours at RT with shaking. Following PBST washes the sulfo-tagged polyGP detection antibody (polyGP antibody [9] conjugated in house with Meso Scale Diagnostics R31AA-1) was diluted 0.5 μg/mL to PBS and added to each well. The plate was incubated with shaking at RT for 1 hour. After washing with PBST, MSD Gold Read Buffer A (Meso Scale Diagnostics R92TG-2) was added to each well and the plate was read with the MESO QuickPlex SQ MSD machine. The polyGP signal value for each sample was then calculated with MSD Discovery Workbench 4.0 Analysis software and the average of each sample triplicate plotted.

### RNA isolation, cDNA synthesis, and RT-qPCR

Total RNA from brain and spinal cord was isolated using the standard TriZol protocol followed by purification using the Direct-zol RNA Miniprep or Microprep kit (Zymo Research R2052). The concentration of RNA samples was determined using a NanoDrop One (Thermo Fisher Scientific). 1μg RNA was used for cDNA synthesis using the High Capacity cDNA Reverse Transcription Kit (Applied Biosystems 4368814). qRT-PCR reactions were performed using TaqMan Fast Advanced Master Mix and an Applied Biosystems StepOnePlus Real Time PCR machine (Applied Biosystems) using commercially available primer-probe Taqman sets (Thermo Fisher Scientific) with Taqman Fast Advanced Master Mix (ThermoFisher Scientific 4444557) or primers and PowerUp Sybr Green Master Mix (Applied Biosystems A25741). **Table 1** contains a list of all Taqman probes and primers used within the study. 10-20 ng of cDNA was used for each reaction and mRNA abundance was measured relative to *Gapdh* mRNA.

**Table 1.**
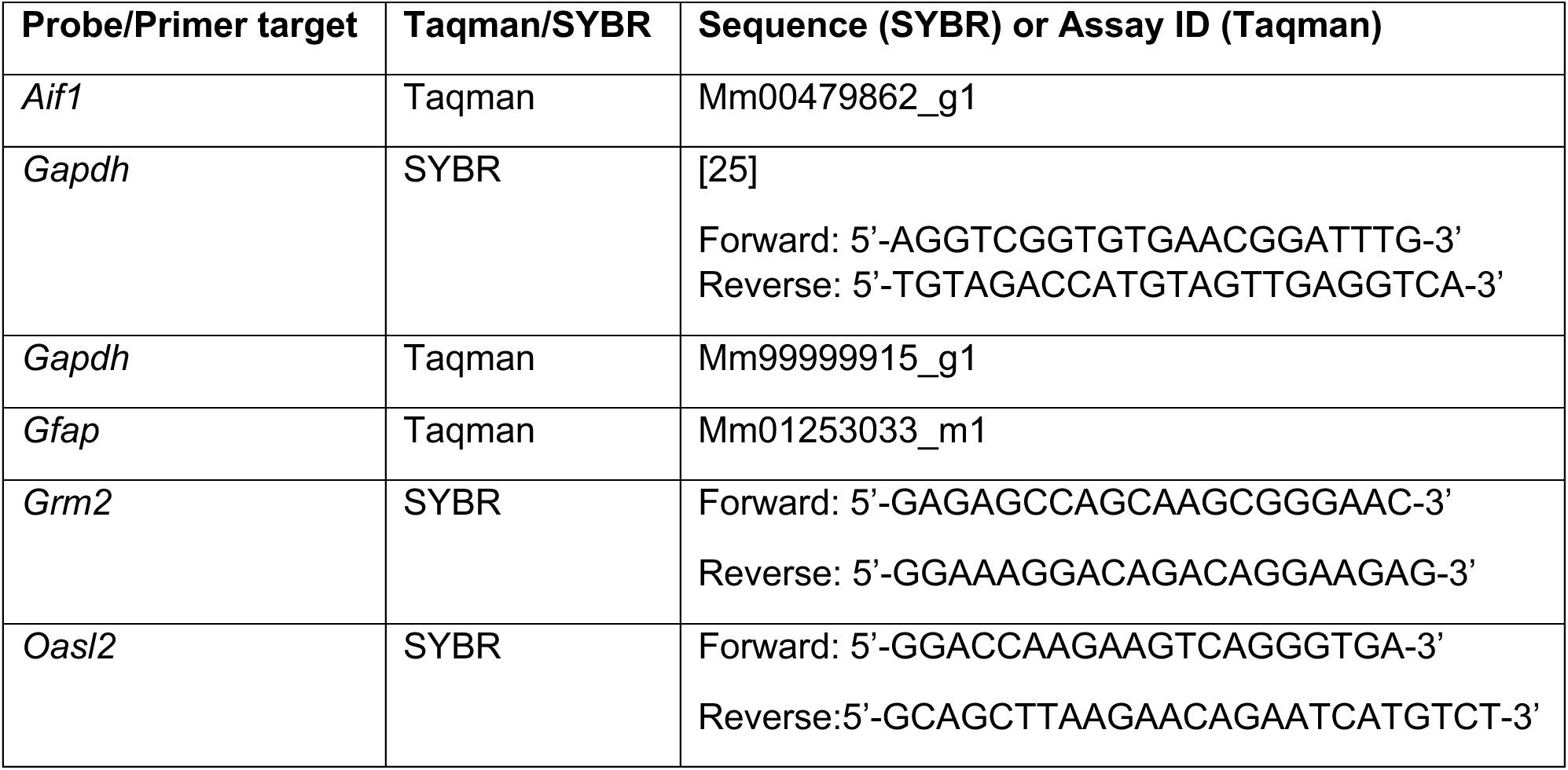
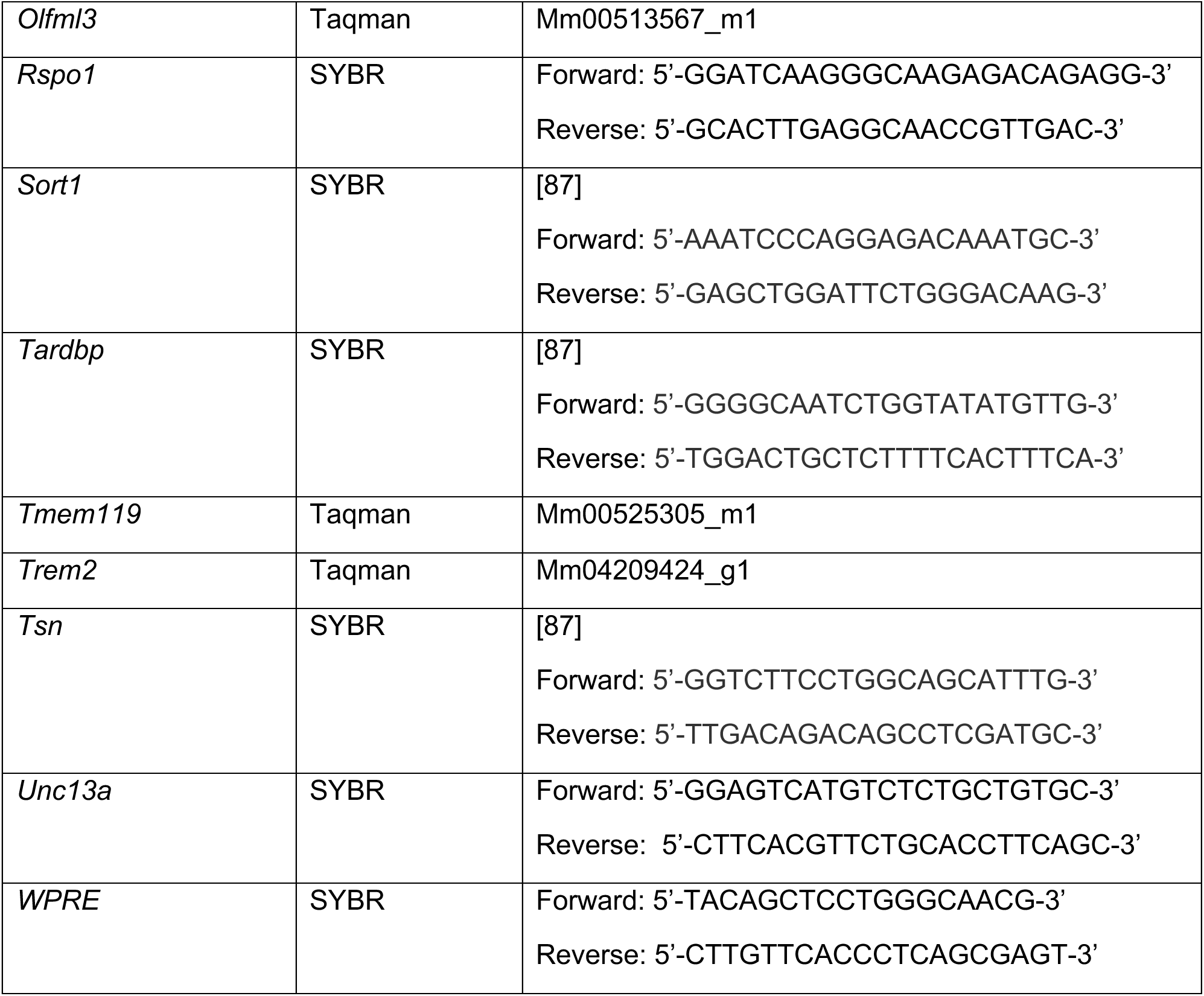
List of qRT-PCR primers and probes.

Technical duplicates or triplicates were utilized within each qRT-PCR plate. A negative control (NTC) was used for each probe to ensure the absence of detected background signal. qRT-PCR results are reported as delta C_T_ values (target C_T_ value – endogenous control C_T_ value) or as Fold Change using the Comparative C_T_ method [43].

### RNA Fluorescence In Situ Hybridization

Formalin-fixed paraffin-embedded (FFPE) sections were prepared for RNA Fluorescence *In Situ* Hybridization with RNAscope reagents (Advanced Cell Diagnostics) according to manufacturer’s instructions. In brief, tissue sections were dried at 60°C for 1 hour and then deparaffinized in xylene and rehydrated through a series of ethanol solutions. Next endogenous peroxidase activity was blocked with RNAscope Hydrogen Peroxide solution for 10 minutes at room temperature. Following distilled water washes, antigen retrieval was performed with 1X Target Retrieval Reagent at 99°C in a steamer for 15 minutes. Slides were next exposed to 100% ethanol for 3 minutes and dried at room temperature. RNAscope Protease Plus solution was added to each section and incubated at 40°C for 30 minutes and washed with distilled water. A probe designed to detect the 5’ and 3’ UTR of the virally expressed RNA (Advanced Cell Diagnostics, RNAscope^TM^ Probe-hC9orf72-BGH-PolyA-C1, 1240972-C1) was added and incubated at 40°C for 2 hours and subsequently the RNAscope Multiplex Fluorescence v2 assay reagents were used as directed. Lastly, sections were incubated with DAPI for 30 seconds at room temperature, mounted with ProLong Gold (ThermoFisher Scientific P36934), and dried overnight.

### Immunofluorescence

FFPE sections were deparaffinized in xylene and rehydrated through a series of ethanol solutions. Antigen retrieval was performed in 10 mM sodium citrate buffer, pH 6.0 for 60 minutes in a steamer and then allowed to cool for 10 minutes. Following washing with deionized water and PBS, the tissue was permeabilized with 0.2% Triton X-100 in PBS for 10 minutes at room temperature. The sections were then washed with PBS with 0.05% Tween (PBST) and blocked with 10% Normal Goat serum containing 0.1 - 0.05% Tween for 1 hour at room temperature. Sections were immunostained with primary antibodies (refer to Table 2) diluted in blocking buffer overnight at 4°C and were subsequently washed with PBST (PBS with 0.05% Tween). Goat secondary antibodies diluted in blocking solution were incubated at room temperature for 1 hour. The sections were thoroughly washed with PBST and mounted on a coverslip with Prolong Gold with DAPI mounting solution (ThermoFisher Scientific P36931).

**Table 2.**
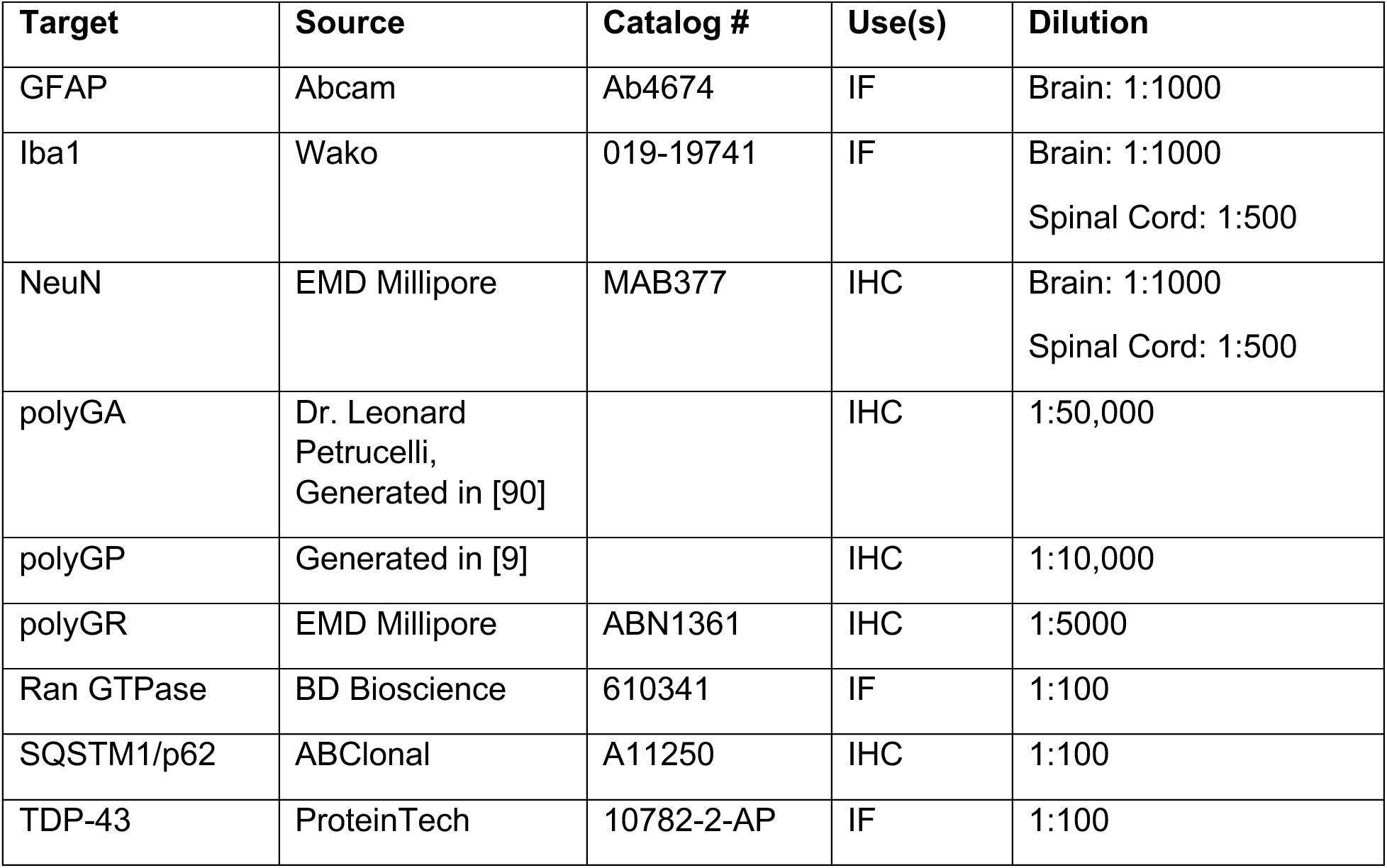
List of antibodies for tissue staining.

| **Target** | **Source** | **Catalog #** | **Use(s)** | **Dilution** |
| --- | --- | --- | --- | --- |
| GFAP | Abcam | Ab4674 | IF | Brain: 1:1000 |
| Iba1 | Wako | 019-19741 | IF | Brain: 1:1000<br>Spinal Cord: 1:500 |
| NeuN | EMD Millipore | MAB377 | IHC | Brain: 1:1000<br>Spinal Cord: 1:500 |
| polyGA | Dr. Leonard Petrucelli,<br>Generated in [90] |  | IHC | 1:50,000 |
| polyGP | Generated in [9] |  | IHC | 1:10,000 |
| polyGR | EMD Millipore | ABN1361 | IHC | 1:5000 |
| Ran GTPase | BD Bioscience | 610341 | IF | 1:100 |
| SQSTM1/p62 | ABClonal | A11250 | IHC | 1:100 |
| TDP-43 | ProteinTech | 10782-2-AP | IF | 1:100 |

### Immunohistochemistry

FFPE sections were deparaffinized in xylene and rehydrated through a series of ethanol solutions. Antigen retrieval was performed in either distilled water (for polyGP, GA, GR staining) or in 10 mM sodium citrate buffer, pH 6.0 (for p62 and NeuN stainings) for 30 minutes. Tissues were immunostained with the following primary antibodies poly(GP), poly(GR), poly(GA), SQSTM1/p62, and NeuN. DAKO Envision+HRP polymer kits (K4003 and K4001) were used and the reaction visualized using ImmPACT® VIP Substrate Kit (Vector Laboratories SK-4605). Sections were then counterstained with Hematoxylin QS Counterstain (Vector Laboratories H-3404-100) and mounted with Aqua-Poly/Mount (Polysciences 18606-20). Slides were either imaged as 20x magnification tiles or individual 63x magnification images using Zeiss Axio Imager.

### NfL and GFAP assessment

Neurofilament light chain (NfL) and GFAP from plasma and CSF samples were assessed using the Quanterix® Neuro 2-Plex B assay (Quanterix 103520). Following the manufacturer’s instructions, samples were blinded and run-in duplicate, along with two quality controls (high-concentration and low-concentration quality control) on each plate per run using the Quanterix SR-X™Analyzer® platform version 1.1. Plasma samples (60 μl plasma into 180 μl of sample diluent) were run using a 4-fold dilution while CSF samples (2.5 μl into 247.5 μl of sample diluent) were run using a 100-fold dilution and results were compensated for the dilution factor. NfL and GFAP concentrations (pg/mL) were calculated using a standard curve (Four parameter logistic (4PL) curve) provided by Quanterix® (Lot 503560).

## Results

### The C9ORF72 (G4C2)66 viral construct is expressed in brain and spinal cord

We utilized a previously generated [10, 75] adeno-associated viral vector (AAV2/9) to drive expression of either 2 (control) or 66 G_4_C_2_ repeats within the intronic region of the chromosome 9 open reading frame 72 (*C9ORF72*) gene. C57BL/6J postnatal day 0 (p0) pups were administered either AAV2/9-*C9ORF72* (G_4_C_2_)_2_ or AAV2/9-*C9ORF72* (G_4_C_2_)_66_ viral particles via intracerebroventricular (ICV) injection (**Supplemental Figure 1A**). To assess the expression of the viral constructs within the CNS, specifically the brain and spinal cord, qRT-PCR was performed on isolated total RNA samples at 180 days following ICV injection (dpi). We observed viral expression in both the (G_4_C_2_)_2_ and (G_4_C_2_)_66_ groups but not the PBS injected (sham) group as expected in the tissues (**Figure 1A, 1C**). The *C9ORF72* HRE leads to the expression of repeat-containing transcripts and formation of RNA foci in C9 ALS/FTD patients [17, 52]. RNA foci were only detectable in the brain and spinal cord of 66R animals and not observed in sham or (G_4_C_2_)_2_ animals (**Figure 1B, 1D**). In the (G_4_C_2_)_66_ animals, foci were found across all layers of the cortex, with elevated expression in layers II and III. We also detected a significant burden of foci within the hippocampus, highest in the Cornu Ammonis (CA) regions, and in the thalamus (**Figure 1B**). Within the spinal cord, we observed the highest level of RNA foci in the ventral horn of cervical sections from (G_4_C_2_)_66_ animals, while no foci were detectable in sham or (G_4_C_2_)_2_ animals (**Figure 1D**). We did not observe RNA foci in lumbar spinal cord sections from sham, (G_4_C_2_)_66_, or (G_4_C_2_)_66_ animals (**Figure 1D**). In **Supplemental Figure 1B**, we have illustrated the collective distribution of the RNA foci that we observed in the brain and spinal cord of (G_4_C_2_)_66_ animals.

**Figure 1.**
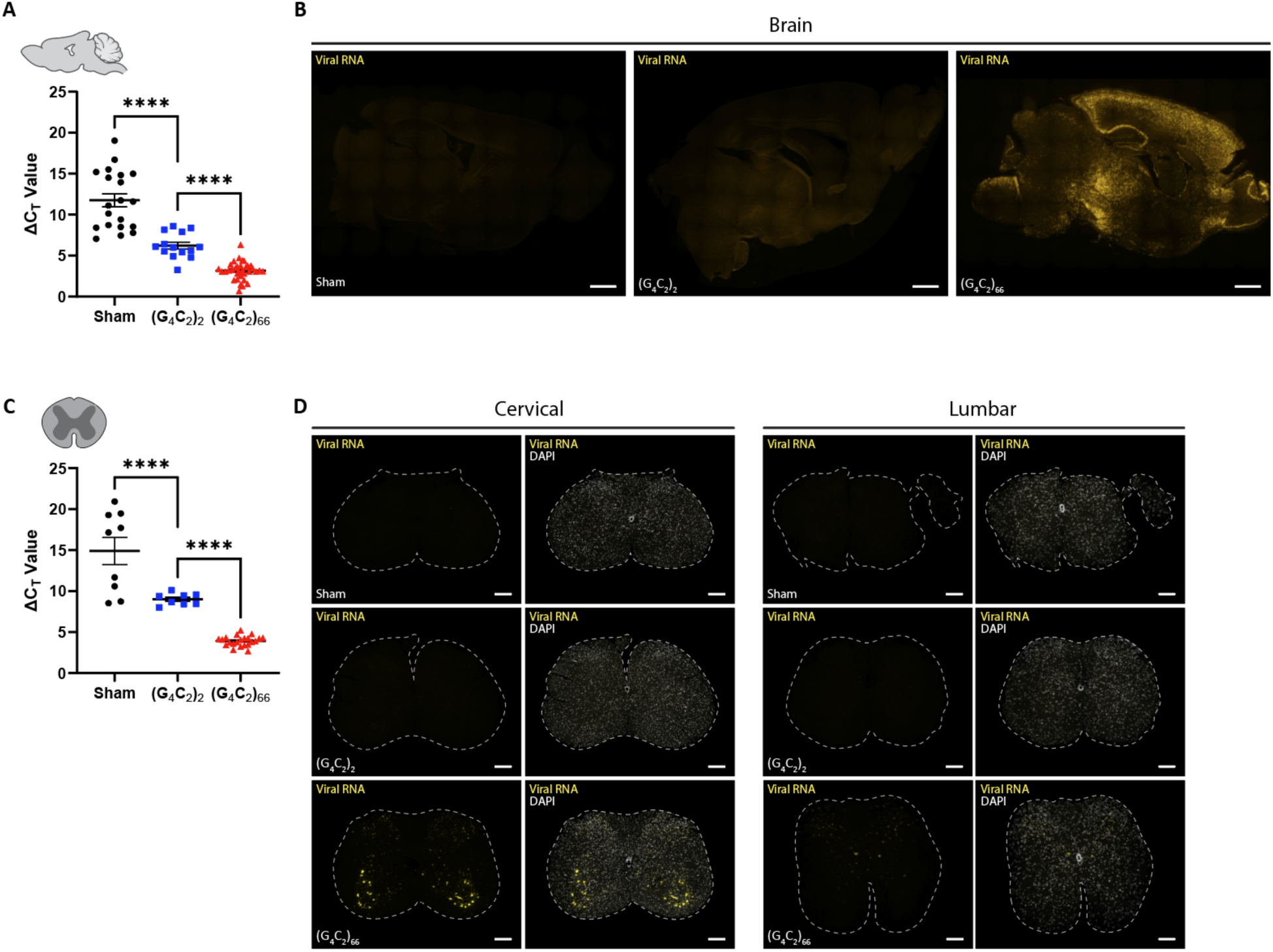
The (G_4_C_2_)_66_ viral construct is expressed in brain and spinal cord following intraventricular delivery. Isolated RNA samples from brain (sham n=20, (G_4_C_2_)_2_ n=14, (G_4_C_2_)_66_ n=34) **(A)** and thoracic spinal cord (sham n=9, (G_4_C_2_)_2_ n=8, (G_4_C_2_)_66_ n=22), error bars = SEM **(C)** were analyzed by qRT-PCR for the expression of the WPRE region of the viral construct. Outliers were identified with the ROUT test (Q=1%) and removed from the dataset. One-way ANOVA analysis of brain (p <0.0001) and spinal cord (p<0.0001) was performed, followed by Tukey’s multiple comparisons test (brain: sham vs. (G_4_C_2_)_2_ p<0.0001, (G_4_C_2_)_2_ vs. (G_4_C_2_)_66_ p<0.0001; spinal cord: sham vs. (G_4_C_2_)_2_ p<0.0001, (G_4_C_2_)_2_ vs. (G_4_C_2_)_66_ p<0.0001), error bars = SEM. Formalin-fixed paraffin-embedded sections from brain and spinal cord sections were analyzed for WPRE expression by RNAScope. Representative images in brain and spinal cord are shown in **(B)** and **(D)**, respectively (sham n=3, (G_4_C_2_)_2_ n=3, (G_4_C_2_)_66_ n=6). Scale bars 1000 µm in (B) and 200 µm in (D).

### C9ORF72 (G4C2)66 mice display DPR expression in the brain and spinal cord

The *C9ORF72* HRE associated DPR peptides are produced by RAN translation [3, 38, 51, 56] from sense and antisense RNA. When we examined the tissue homogenates ny polyGP MSD, only (G_4_C_2_)_66_ mice had a robust polyGP signal in both the brain and spinal cord (**Figure 2A**). Importantly, the significant increase in polyGP signal detected in the brain of (G_4_C_2_)_66_ mice was completely abolished with the delivery of a *C9ORF72* anti-sense oligonucleotide (C9-ASO) at 90 dpi via intracerebroventricular (ICV) injection, with no statistical difference found between the sham and (G_4_C_2_)_66_ + C9-ASO groups in the brain (**Figure 2A**). We also measured polyGP signal in thoracic spinal cord samples and found a significant increase in polyGP signal in the (G_4_C_2_)_66_ mice (**Figure 2A**). We observed approximately 10-fold less polyGP signal in spinal cord compared to brain polyGP signal in (G_4_C_2_)_66_ animals—indicative of a greater polyGP burden in the brain than the spinal cord of (G_4_C_2_)_66_ animals (**Figure 2A**)—consistent with the reduced viral loads observed (**Figure 1C**). Additionally, to gain insight into the expression and location of polyGP, polyGR, and polyGA species *in situ*, we performed immunohistochemistry on brain and spinal cord tissues. In the (G_4_C_2_)_66_ transduced animals we observed robust polyGP signal within the nucleus of cells within the cortex, thalamus, and hippocampus, but not in (G_4_C_2_)_2_ and sham animals (**Figure 2B-C, Supplemental Figure 2A-B**). For both polyGR and polyGA we observed smaller, aggregates perinuclear or within the nucleus of cortical, thalamic, and hippocampal cells of (G_4_C_2_)_66_ animals (**Figure 2D-G, Supplemental Figure 2A-B**). The DPR positive cells within each brain region was quantified; the expression ranged from an average of approximately 30 cells per millimeter squared (mm^2^) in the cortex for polyGA up to 150 cells per mm^2^ in the thalamus for polyGR. A significant increase in DPR positive cells in the (G_4_C_2_)_66_ group was found when compared to controls in all cases, except for polyGP positive cells in the cortex where there was significant variability between animals (**Figure 2C**, **2E**, **2G**). In **Supplemental Figure 1C**, we have illustrated the collective distribution of the DPR species that we observed in the brain of (G_4_C_2_)_66_ animals.

**Figure 2.**
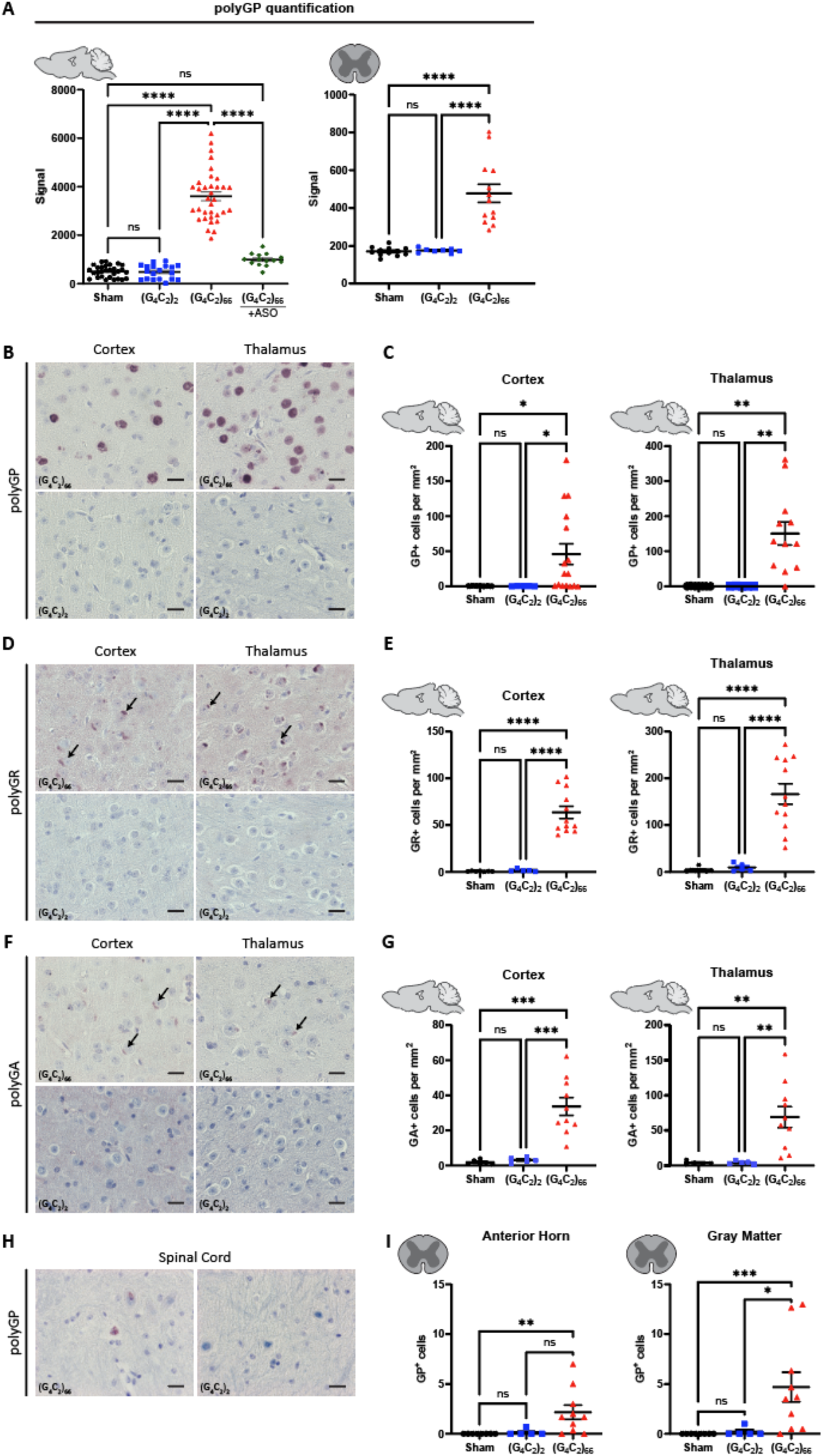
(G_4_C_2_)_66_ mice have DPR expression in the brain and spinal cord. **(A)** Brain (sham n=29, (G_4_C_2_)_2_ n=19, (G_4_C_2_)_66_ n=34, (G_4_C_2_)_66_ +C9-ASO n=14) and thoracic spinal cord (sham n=12, (G_4_C_2_)_2_ n=8, (G_4_C_2_)_66_ n=13) tissue lysates were analyzed by Meso Scale Discovery assay for the expression of polyGP dipeptide repeat. One-way ANOVA analysis of brain (p <0.0001) and spinal cord (p<0.0001) was performed, followed by Tukey’s multiple comparisons test (brain: sham vs. (G_4_C_2_)_2_ p=0.9996, sham vs. (G_4_C_2_)_66_ p<0.0001, (G_4_C_2_)_2_ vs. (G_4_C_2_)_66_ p<0.0001, sham vs. (G_4_C_2_)_66_ +C9-ASO p=0.1123, (G_4_C_2_)_66_ vs. (G_4_C_2_)_66_ + C9-ASO p<0.0001; spinal cord: sham vs. (G_4_C_2_)_2_ p=0.9958, sham vs. (G_4_C_2_)_66_ p<0.0001, (G_4_C_2_)_2_ vs. (G_4_C_2_)_66_ p<0.001, error bars = SEM. **(B)** Representative images of brain sections stained for polyGP. Scale bars 20 µm. **(C)** Quantification of polyGP positive cells in cortex and thalamus region (sham n=7, (G_4_C_2_)_2_ n=5, (G_4_C_2_)_66_ n=16 for cortex, 12 for hippocampus and thalamus). One-way Welch ANOVA analysis (Welch, 1947) of polyGP positive cell number was performed (p=0.0021 for cortex and p=0.0002 for thalamus) followed by Tukey’s multiple comparison test. Cortex: sham vs (G_4_C_2_)_2_ p=0.4842, sham vs (G_4_C_2_)_66_ p=0.0220 and (G_4_C_2_)_2_ vs (G_4_C_2_)_66_ p=0.0215, Thalamus: sham vs (G_4_C_2_)_2_ p>0.9999, sham vs (G_4_C_2_)_66_ p=0.0023 and (G_4_C_2_)_2_ vs (G_4_C_2_)_66_ p=0.0023, error bars = SEM. **(D)** Representative images of brain sections stained for polyGR. Scale bars 20 µm. **(E)** Quantification of polyGR positive cells in cortex and thalamus region (sham n=7, (G_4_C_2_)_2_ n=5, (G_4_C_2_)_66_ n=12). One-way Welch ANOVA analysis of polyGR positive cell number was performed (p<0.0001 for cortex and thalamus) followed by Tukey’s multiple comparison test. Cortex: sham vs (G_4_C_2_)_2_ p=0.4769, sham vs (G_4_C_2_)_66_ p<0.0001 and (G_4_C_2_)_2_ vs (G_4_C_2_)_66_ p<0.0001, Thalamus: sham vs (G_4_C_2_)_2_ p=0.5929, sham vs (G_4_C_2_)_66_ p<0.0001 and (G_4_C_2_)_2_ vs (G_4_C_2_)_66_ p<0.0001, error bars = SEM. **(F)** Representative images of brain sections stained for polyGA. Scale bars 20µm. **(G)** Quantification of polyGA positive cells in cortex and thalamus region (sham n=6 for cortex, 5 for thalamus and hippocampus, (G_4_C_2_)_2_ n=5, (G_4_C_2_)_66_ n=10 for cortex and thalamus, 9 for hippocampus). One-way Welch ANOVA analysis of polyGA positive cell number was performed (p<0.0001 for cortex and p=0.0006 for thalamus) followed by Tukey’s multiple comparison test. Cortex: sham vs (G_4_C_2_)_2_ p=0.3688, sham vs (G_4_C_2_)_66_ p=0.0004 and (G_4_C_2_)_2_ vs (G_4_C_2_)_66_ p=0.0006, Thalamus: sham vs (G_4_C_2_)_2_ p=0.9980, sham vs (G_4_C_2_)_66_ p=0.0056 and (G_4_C_2_)_2_ vs (G_4_C_2_)_66_ p=0.0057, error bars = SEM. **(H)** Representative images of cervical spinal cord sections stained against polyGP. Scale bars 20 µm. **(I)** Quantification of cervical spinal cord sections stained against polyGP (sham n=8, (G_4_C_2_)_2_ n=5, (G_4_C_2_)_66_ n=10). A Kruskal-Wallis test (Kruskal, Wallis 1952) on the cervical spinal cord gray matter (p=0.0006) and anterior horn (p=0.0018) was performed, followed by Dunn’s multiple comparisons test (Dunn, 1964) (gray matter: sham vs. (G_4_C_2_)_2_ p>0.9999, sham vs. (G_4_C_2_)_66_ p=0.0008, (G_4_C_2_)_2_ vs. (G_4_C_2_)_66_ p=0.0266; anterior horn: sham vs. (G_4_C_2_)_2_ p>0.9999, sham vs. (G_4_C_2_)_66_ p=0.0024, (G_4_C_2_)_2_ vs. (G_4_C_2_)_66_ p=0.0512), error bars = SEM.

In the spinal cord of (G_4_C_2_)_66_ animals we found a less robust DPR burden than in the brain, in concordance with our polyGP MSD and RNAscope results (**Figure 2A**; **Figure 1C**). Specifically, we observed a significant increase in polyGP positive cells in the cervical spinal cord sections (gray matter and anterior horn), but on average we only observed between 3-5 positive cells per section (**Figure 2H-I**). In lumbar spinal cord sections, we observed a slight but significant increase in polyGP positive cells in gray matter but not in the anterior horn (**Supplemental Figure 2C**). There was no detectable expression of polyGR or polyGA positive cells in the spinal cord of (G_4_C_2_)_66_ animals by IHC (data not shown).

### TDP-43 and RanGTPase are functional in the C9ORF72 (G4C2)66 mouse model

TDP-43 dysfunction is a hallmark feature in ALS, and several mouse models of ALS have recapitulated aggregated TDP-43 pathology *in situ* [45, 63]. We assessed the *C9ORF72* (G_4_C_2_)_66_ mouse model for indicators of TDP-43 dysfunction. Loss of nuclear TDP-43 is an important event in ALS pathogenesis [65], thus we utilized immunostaining of total TDP-43 expression and localization as a readout for TDP-43 dysfunction. When we compared the expression of TDP-43 in neurons in the motor cortex of (G_4_C_2_)_66_ animals to sham and (G_4_C_2_)_2_ animals we did not observe any gross change in TDP-43 distribution or nuclear versus cytoplasmic (N/C) ratio (**Figure 3A-B**). Moreover, we evaluated localization of TDP-43 in cells with significant DPR burden in various brain regions in (G_4_C_2_)_66_ mice and did not observe mislocalization of TDP-43 in DPR positive cells (**Supplemental Figure 3**). TDP-43 performs multiple functions, such as transcriptional repression, splicing, and translational regulation, that are disrupted in ALS [40, 76, 80]. The consequence of TDP-43 mislocalization can be evaluated by interrogating TDP-43 RNA targets [6, 33, 40, 46, 64]. We quantified the expression of several previously documented TDP-43 mRNA targets [86] in the brain and spinal cord of (G_4_C_2_)_66_ animals. When compared to control groups, we found that (G_4_C_2_)_66_ animals had no significant change in the expression of *Grm2*, *Oasl1*, *Rspo1*, *Sort1*, *Tardbp*, *Tsn*, and *Unc13a* (**Figure 3C-D**). Finally, we stained for RanGTPase as an indicator of nucleocytoplasmic transport (NCT) function in (G_4_C_2_)_66_ animals, as loss of a steep nuclear-to-cytoplasmic RanGTPase gradient may lie upstream of TDP-43 dysfunction in ALS [13]. We observed a slight but significant increase in the RanGTPase N/C ratio in (G_4_C_2_)_66_ compared to (G_4_C_2_)_2_ animals, but not when (G_4_C_2_)_66_ animals were compared to sham animals (**Figure 3E-F**). Thus, it is unlikely that neither TDP-43 function or nucleocytoplasmic transport is significantly altered in the neurons of (G_4_C_2_)_66_ mice.

**Figure 3.**
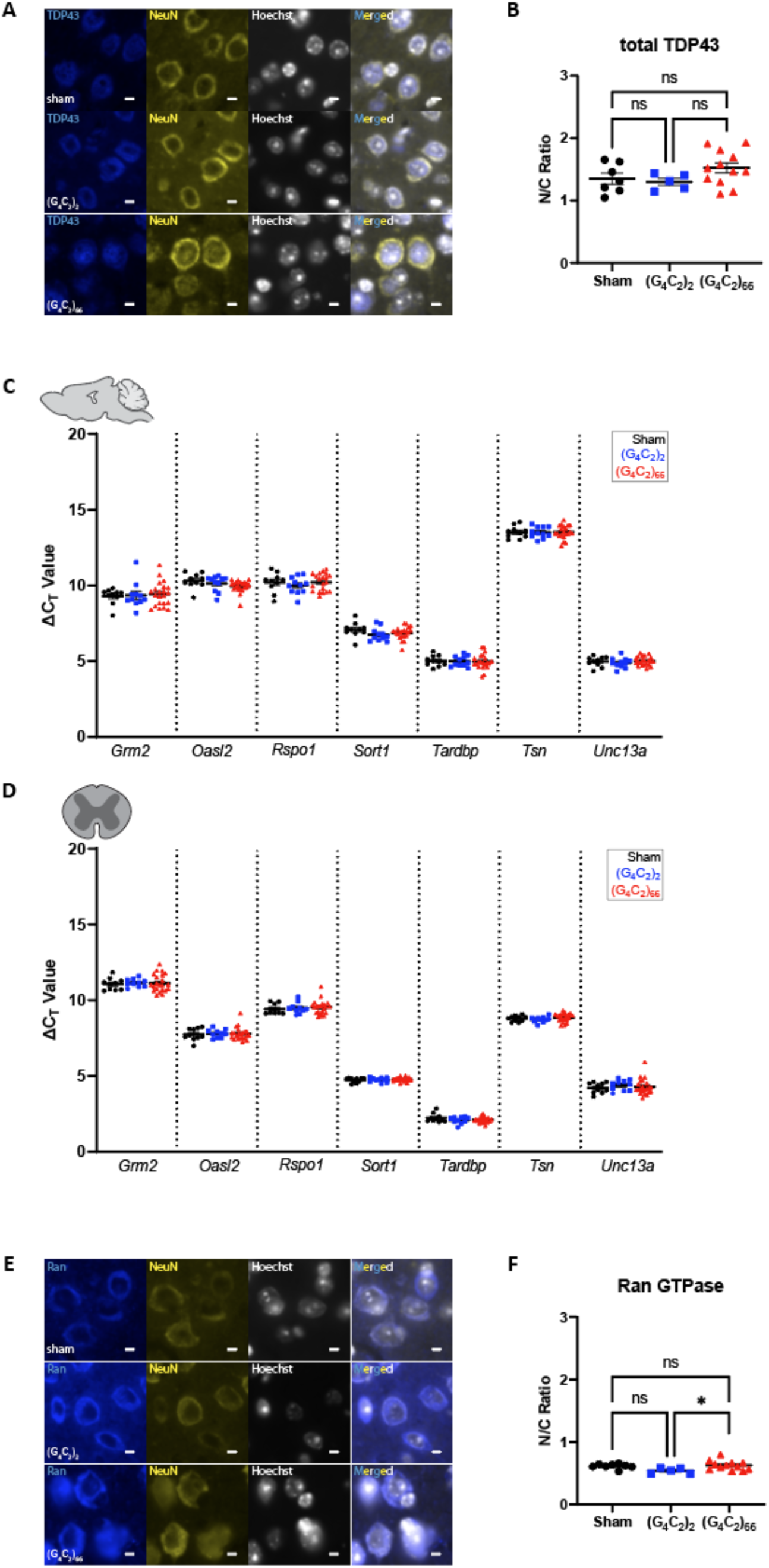
No evidence for loss of TDP-43 or RanGTPase function in the (G_4_C_2_)_66_ mouse model. **(A)** Representative images of immunohistochemistry analysis of TDP-43 expression within the motor cortex tissue. Scale bars 5 µm. **(B)** Quantification of nuclear to cytoplasmic ratio of TDP-43 (sham n=7, (G_4_C_2_)_2_ n=5, (G_4_C_2_)_66_ n=12). One-way ANOVA: p=0.1597, Tukey’s multiple comparisons test: sham vs. (G_4_C_2_)_2_ p=0.9297, sham vs. (G_4_C_2_)_66_ p=0.3097, (G_4_C_2_)_2_ vs. (G_4_C_2_)_66_ p=0.2129. The expression of several TDP-43 mRNA targets was analyzed by qRT-PCR in samples from **(C)** brain (sham n=10, (G_4_C_2_)_2_ n=11, (G_4_C_2_)_66_ n=21) and **(D)** thoracic spinal cord (sham n=10, (G_4_C_2_)_2_ n=10, (G_4_C_2_)_66_ n=22) tissue. Outliers were identified with the ROUT test (Q=1%) and removed from the dataset. A one-way ANOVA was performed for each gene independently and no significant statistical difference was found in brain (*Grm2*: p=0.8822; *Oasl2*: p=0.1082; *Rspo1*: p=0.5920; *Sort1*: p=0.5175; *Tardbp*: p=0.9677; *Tsn*: p=0.9973; *Unc13a*: p=0.5271) or spinal cord (*Grm2*: p=0.9061; *Oasl2*: p=0.9364; *Rspo1*: p=0.7679; *Sort1*: p=0.6396; *Tardbp*: p=0.2842; *Tsn*: p=0.3036; *Unc13a*: p=0.6663), error bars = SEM. **(E)** Representative images of immunohistochemistry analysis of RanGTPase analysis within the motor cortex tissue. Scale bars 5 µm. **(F)** Quantification of nuclear to cytoplasmic ratio of RanGTPase (sham n=8, (G_4_C_2_)_2_ n=5, (G_4_C_2_)_66_ n=12). One-way ANOVA: p=0.0401, Tukey’s multiple comparisons test: sham vs. (G_4_C_2_)_2_ p=0.1101, sham vs. (G_4_C_2_)_66_ p=0.8696, (G_4_C_2_)_2_ vs. (G_4_C_2_)_66_ p=0.0339, error bars = SEM.

### Lack of Neuronal Degeneration but presence of P62-positive inclusions in the brain of C9ORF72 (G4C2)66 mice

Disruptions in protein and RNA homeostasis as a result of an aberrant stress response have been implicated in the pathogenesis of C9 ALS [41]. To evaluate the *C9ORF72* (G_4_C_2_)_66_ repeat expansion model for disturbances in protein homeostasis, we immunostained brain tissue for p62/Sqstm1, a protein that aggregates and associates with stress granules. We found p62-positive inclusions throughout the brain, including the cortex, hippocampus, and thalamus of 6-month old (G_4_C_2_)_66_ animals (**Figure 4A**). The number of p62-positive inclusions in all these brain regions was significantly increased in comparison to both sham and (G_4_C_2_)_2_ animals (**Figure 4B**). These data clearly indicate that the (G_4_C_2_)_66_ repeat expansion leads to aberrant protein accumulation in the AAV-(G_4_C_2_)_66_ mouse model.

**Figure 4.**
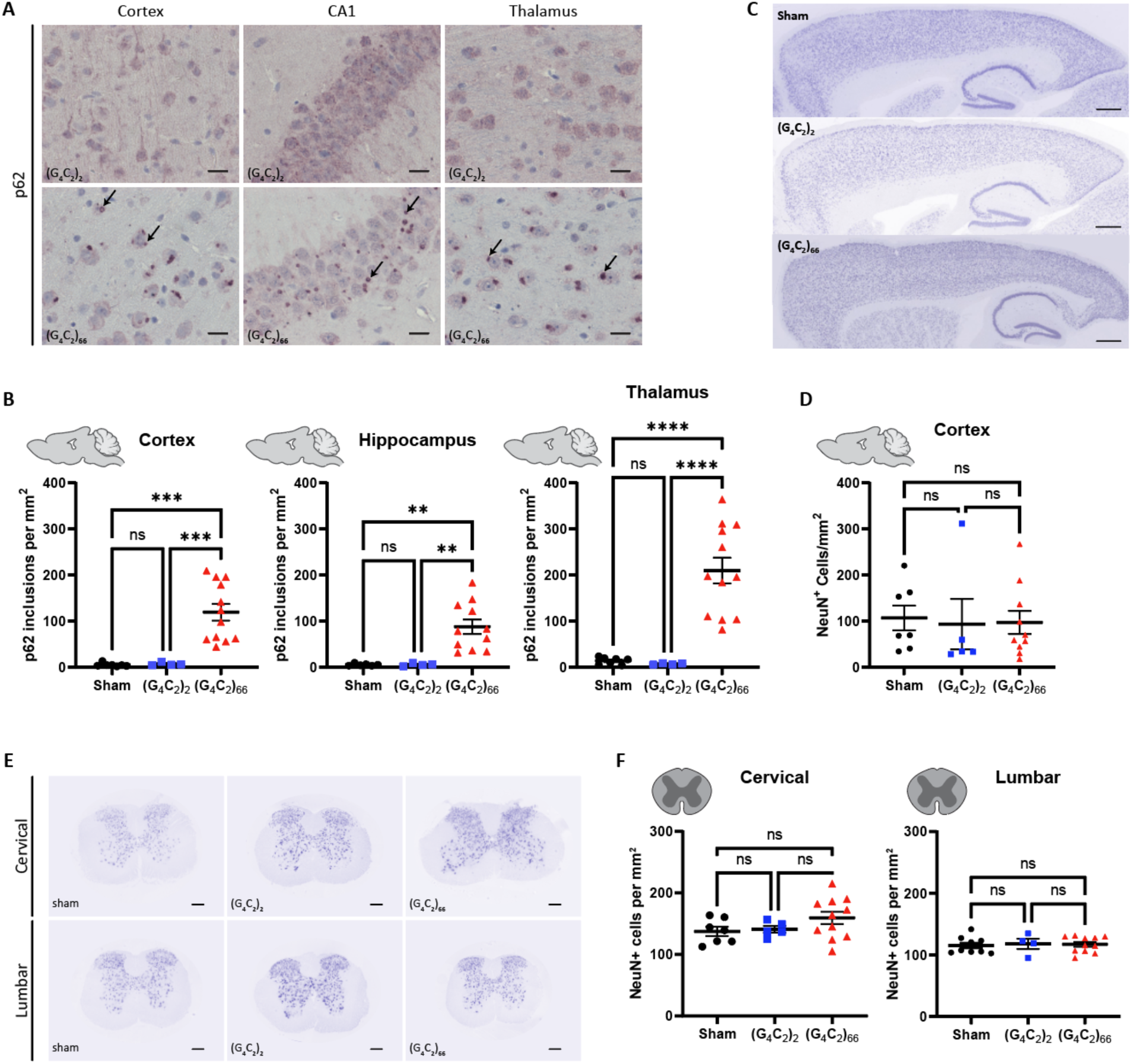
Analysis of p62 inclusions and neuronal count in the (G_4_C_2_)_66_ mouse model. **(A)** Representative images of brain sections stained against p62. As pointed by the arrows, p62 positive inclusions are mostly seen in 6-month old (G_4_C_2_)_66_ mice and not in sham or (G_4_C_2_)_2_ mice. Scale bars 20 µm. **(B)** Quantification of p62 inclusions in cortex, thalamus, and hippocampus region (sham n=7 for thalamus and cortex, (G_4_C_2_)_2_ n=4, (G_4_C_2_)_66_ n=12 for cortex and thalamus, 11 for hippocampus). One-way Welch ANOVA analysis of p62 inclusions was performed (p<0.0001 for cortex, thalamus, and hippocampus) followed by Tukey’s multiple comparison test. Cortex: sham vs (G_4_C_2_)_2_ p=0.7421, sham vs (G_4_C_2_)_66_ p=0.0002 and (G_4_C_2_)_2_ vs (G_4_C_2_)_66_ p=0.0002, Thalamus: sham vs (G_4_C_2_)_2_ p=0.1518, sham vs (G_4_C_2_)_66_ p<0.0001 and (G_4_C_2_)_2_ vs (G_4_C_2_)_66_ p<0.0001, Hippocampus: sham vs (G_4_C_2_)_2_ p=0.9997, sham vs (G_4_C_2_)_66_ p=0.0011 and (G_4_C_2_)_2_ vs (G_4_C_2_)_66_ p=0.0011, error bars = SEM. **(C)** Representative images of brain sections from 6-month old animals stained against NeuN. **(D)** Quantification of NeuN positive cells within the cortex (sham n=7, (G_4_C_2_)_2_ n=5, (G_4_C_2_)_66_ n=10). Scale bars 500 µm. One-way ANOVA: p=0.9601, Tukey’s multiple comparisons test: sham vs. (G_4_C_2_)_2_ p=0.9631, sham vs. (G_4_C_2_)_66_ p=0.9714, (G_4_C_2_)_2_ vs. (G_4_C_2_)_66_ p=0.9971, error bars = SEM. **(E)** Representative cervical and lumbar spinal cord sections stained against NeuN. Scale bars 200 µm. **(F)** Quantification of NeuN positive cells within the cervical (sham n=7, (G_4_C_2_)_2_ n=5, (G_4_C_2_)_66_ n=11) and lumbar spinal cord (sham n=9, (G_4_C_2_)_2_ n=4, (G_4_C_2_)_66_ n=11). One-way ANOVA analysis of the cervical spinal cord dataset was performed (p=0.2056), followed by Tukey’s multiple comparisons test: sham vs. (G_4_C_2_)_2_ p=0.9723, sham vs. (G_4_C_2_)_66_ p=0.2296, (G_4_C_2_)_2_ vs. (G_4_C_2_)_66_ p=0.4228. One-way ANOVA analysis of the lumbar spinal cord dataset was performed (p=0.9341), followed by Tukey’s multiple comparisons test: sham vs. (G_4_C_2_)_2_ p=0.9448, sham vs. (G_4_C_2_)_66_ p=0.9521, (G_4_C_2_)_2_ vs. (G_4_C_2_)_66_ p=0.9944, error bars = SEM.

The hallmark pathology of ALS is progressive degeneration of both upper and lower motor neurons. We assessed 6-month old mice injected with the (G_4_C_2_)_66_ repeat expansion for neuronal loss by immunostaining brain and spinal cord sections with neuronal marker NeuN. The number of NeuN-positive neurons in the cortex of 7 sham, 5 (G_4_C_2_)_2_ and 10 (G_4_C_2_)_66_ animals was quantified (**Figure 4C-D**). Surprisingly, we found that there was no significant difference in the number of NeuN-positive cells between groups. Next, we quantified the number of NeuN-positive cells in both cervical and lumbar spinal cord sections from control and (G_4_C_2_)_66_ animals, where we also found that there was no difference between groups (**Figure 4E-F**). Collectively, this lack of observed neuronal loss suggests that expression of the (G_4_C_2_)_66_ construct does not cause apparent neurodegeneration in the (G_4_C_2_)_66_ mouse model.

### Mild gliosis is observed in the brain, but not the spinal cord of C9ORF72 (G4C2)66 mice

We assessed the cellular response of astrocytes and microglia to the (G_4_C_2_)_66_ repeat expansion with immunofluorescence and qRT-PCR at 6-month of age. First, we quantified glial fibrillary acidic protein (GFAP) immunoreactivity in brain and spinal cord sections. The percent GFAP-positive area did not significantly differ between sham, (G_4_C_2_)_2_, and (G_4_C_2_)_66_ animals in the cortex or cervical spinal cord tissue (**Figure 5A-B**). Similarly, we quantified the GFAP-positive area in the motor cortex and lumbar spinal cord and found no difference across experimental groups (**Supplemental Figure 4A-B**). We utilized qRT-PCR for increased sensitivity to analyze *Gfap* expression in the brain and thoracic spinal cord tissue. Surprisingly, we found a significant increase in *Gfap* expression in the brain, but not the spinal cord of (G_4_C_2_)_66_ animals in comparison to sham animals (**Figure 6A-B**), suggesting there may be mild astrocyte reactivity in the brains of (G_4_C_2_)_66_ animals.

**Figure 5.**
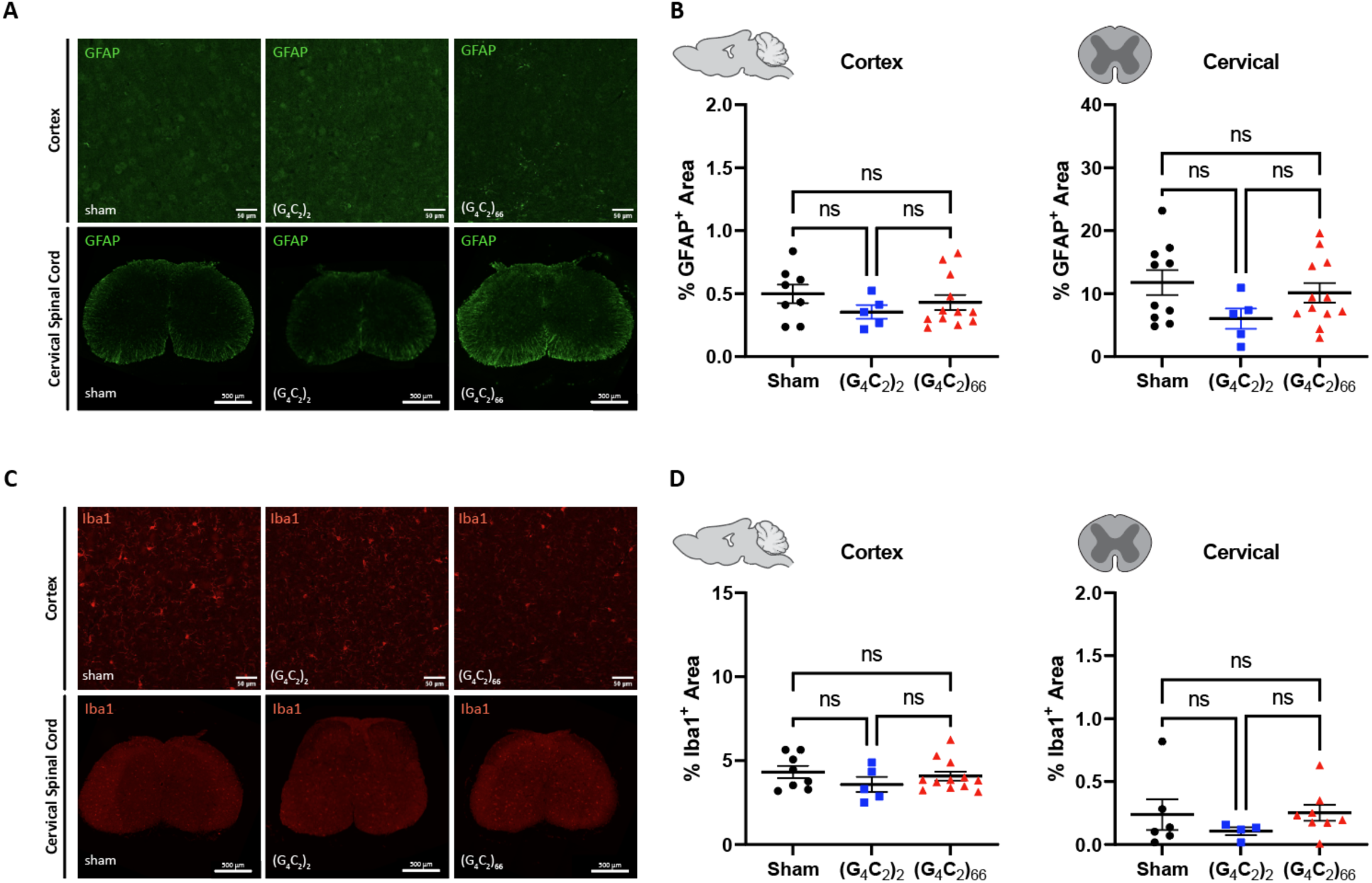
Analysis of gliosis in the 6-month old (G_4_C_2_)_66_ mouse model by immunohistochemistry. **(A)** Representative images of immunohistochemistry analysis of GFAP expression within the cortex and cervical spinal cord. Cortex scale bars 50 µm and cervical spinal cord scale bars 200 µm. **(B)** Quantification of % GFAP positive area within cortex (sham n=8, (G_4_C_2_)_2_ n=5, (G_4_C_2_)_66_ n=12) and cervical spinal cord (sham n=10, (G_4_C_2_)_2_ n=5, (G_4_C_2_)_66_ n=12). One-way ANOVA analysis of the cortex dataset was performed (p=0.4382), followed by Tukey’s multiple comparisons test: sham vs. (G_4_C_2_)_2_ p=0.4110, sham vs. (G_4_C_2_)_66_ p=0.7243, (G_4_C_2_)_2_ vs. (G_4_C_2_)_66_ p=0.7490. One-way ANOVA analysis of the cervical spinal cord dataset was performed (p=0.1798), followed by Tukey’s multiple comparisons test: sham vs. (G_4_C_2_)_2_ p=0.1558, sham vs. (G_4_C_2_)_66_ p=0.7651, (G_4_C_2_)_2_ vs. (G_4_C_2_)_66_ p=0.3528, error bars = SEM. **(C)** Representative images of immunohistochemistry analysis of Iba1 expression within cortex and cervical spinal cord. Cortex scale bars 50 µm and cervical spinal cord scale bars 200 µm. **(D)** Quantification of % Iba1 positive area within cortex (sham n=8, (G_4_C_2_)_2_ n=5, (G_4_C_2_)_66_ n=12) and cervical spinal cord (sham n=6, (G_4_C_2_)_2_ n=4, (G_4_C_2_)_66_ n=8). One-way ANOVA analysis of the cortex dataset was performed (p=0.4234), followed by Tukey’s multiple comparisons test: sham vs. (G_4_C_2_)_2_ p=0.3924, sham vs. (G_4_C_2_)_66_ p=0.8465, (G_4_C_2_)_2_ vs. (G_4_C_2_)_66_ p=0.6124. One-way ANOVA analysis of the cervical spinal cord dataset was performed (p=0.5281), followed by Tukey’s multiple comparisons test: sham vs. (G_4_C_2_)_2_ p=0.6173, sham vs. (G_4_C_2_)_66_ p=0.9922, (G_4_C_2_)_2_ vs. (G_4_C_2_)_66_ p=0.5239, error bars = SEM.

**Figure 6.**
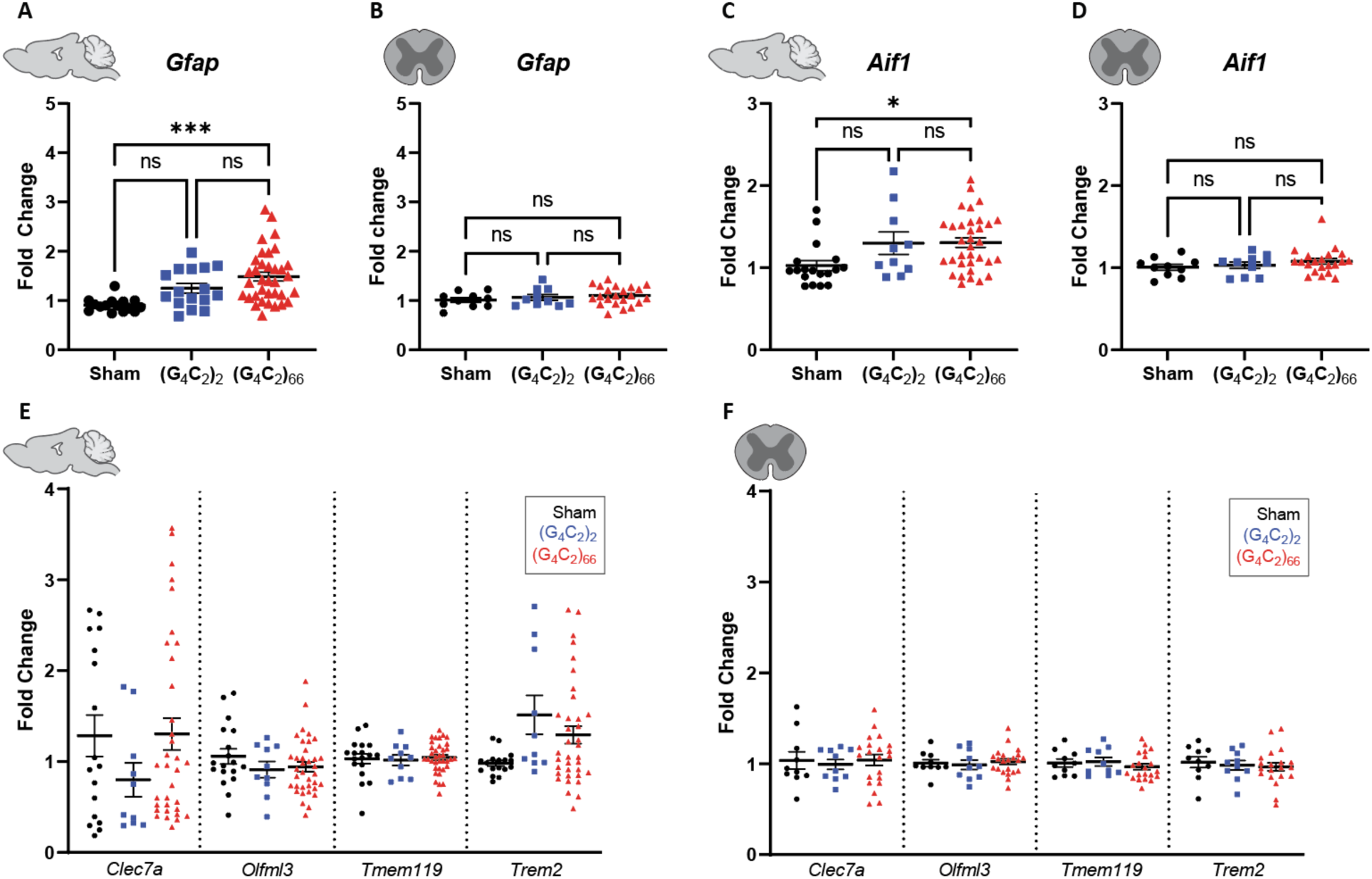
Analysis of gliosis in the (G_4_C_2_)_66_ mouse model by qRT-PCR. Analysis of *Gfap* expression within **(A)** brain (sham n=15, (G_4_C_2_)_2_ n=16, (G_4_C_2_)_66_ n=36) and **(B)** thoracic spinal cord tissue (sham n=10, (G_4_C_2_)_2_ n=10, (G_4_C_2_)_66_ n=22). Outliers were identified with the ROUT test (Q=1%) and removed from the dataset. One-way ANOVA analysis of the brain dataset was performed (p=0.0003), followed by Tukey’s multiple comparisons test: sham vs. (G_4_C_2_)_2_ p=0.0810, sham vs. (G_4_C_2_)_66_ p=0.0002, (G_4_C_2_)_2_ vs. (G_4_C_2_)_66_ p=0.1774. One-way ANOVA analysis of the spinal cord dataset was performed (p=0.8903), followed by Tukey’s multiple comparisons test: sham vs. (G_4_C_2_)_2_ p=0.7816, sham vs. (G_4_C_2_)_66_ p=0.3886, (G_4_C_2_)_2_ vs. (G_4_C_2_)_66_ p=0.8517. Analysis of *Aif1* expression within **(C)** brain (sham n=18, (G_4_C_2_)_2_ n=10, (G_4_C_2_)_66_ n=34) and **(D)** thoracic spinal cord tissue (sham n=10, (G_4_C_2_)_2_ n=10, (G_4_C_2_)_66_ n=21). Outliers were identified with the ROUT test (Q=1%) and removed from the dataset. One-way ANOVA analysis of the brain dataset was performed (p=0.0164), followed by Tukey’s multiple comparisons test: sham vs. (G_4_C_2_)_2_ p=0.1056, sham vs. (G_4_C_2_)_66_ p=0.0156, (G_4_C_2_)_2_ vs. (G_4_C_2_)_66_ p=0.9982. One-way ANOVA analysis of the thoracic spinal cord dataset was performed (p=0.3671), followed by Tukey’s multiple comparisons test: sham vs. (G_4_C_2_)_2_ p=0.9225, sham vs. (G_4_C_2_)_66_ p=0.3785, (G_4_C_2_)_2_ vs. (G_4_C_2_)_66_ p=0.6427. Analysis of *Clec7a*, *Olfml3*, *Tmem119*, and *Trem2* expression within **(E)** brain (sham n=18, (G_4_C_2_)_2_ n=10, (G_4_C_2_)_66_ n=34) and **(F)** thoracic spinal cord tissue (sham n=10, (G_4_C_2_)_2_ n=10, (G_4_C_2_)_66_ n=21). Outliers were identified with the ROUT test (Q=1%) and removed from the dataset. A one-way ANOVA was performed for each gene independently and no significant statistical difference was found in brain (*Clec7a*: p=0.3184; *Olfml3*: p=0.3836; *Tmem119*: p=0.8965; *Trem2*: p=0.1103) or spinal cord (*Clec7a*: p=0.8811; *Olfml3*: p=0.8136; *Tmem119*: p=0.5710; *Trem2*: p=0.7783), error bars = SEM.

To determine if microglia are responsive to the (G_4_C_2_)_66_ repeat expansion, we quantified Iba1 immunoreactivity and morphology in the brain and spinal cord sections from all groups. There was no significant difference in the percent area that was Iba1-positive between 6-month old sham, (G_4_C_2_)_2_, and (G_4_C_2_)_66_ animals across brain and spinal cord sections (**Figure 5C-D**, **Supplemental Figure 4C-D**). Additionally, there was no observable change to microglia morphology across all groups (data not shown). Next, we used qRT-PCR to assess the global expression of microglia markers, including *Aif1*, *Clec7a*, *Olfml3*, *Tmem119*, and *Trem2* (**Figure 6C-F**). Only brain samples from (G_4_C_2_)_66_ animals had a significant increase in *Aif1* expression (**Figure 6C**), suggesting there is a minor microglia response to the expression of the (G_4_C_2_)_66_ repeat expression construct. However, we do not observe significant differences in these glial activation indicators between (G_4_C_2_)_2_ and (G_4_C_2_)_66_ mice, so the minor glial activation may be due to the injection procedure or the virus itself.

Lastly, we assessed the impact of (G_4_C_2_)_66_ expression on systemic and CSF levels of neurofilament light chain (NfL) and GFAP. Despite no detectable loss of neurons and no apparent increase in GFAP immunoreactivity in CNS tissues, we found an increase in NfL concentration in both plasma and CSF compartments in 6-month old (G_4_C_2_)_66_ animals compared to the sham group (**Figure 7A-B**). The GFAP concentration measured in plasma from (G_4_C_2_)_66_ animals were alsoincreased compared to sham, but not in CSF (**Figure 7A-B**).

**Figure 7.**
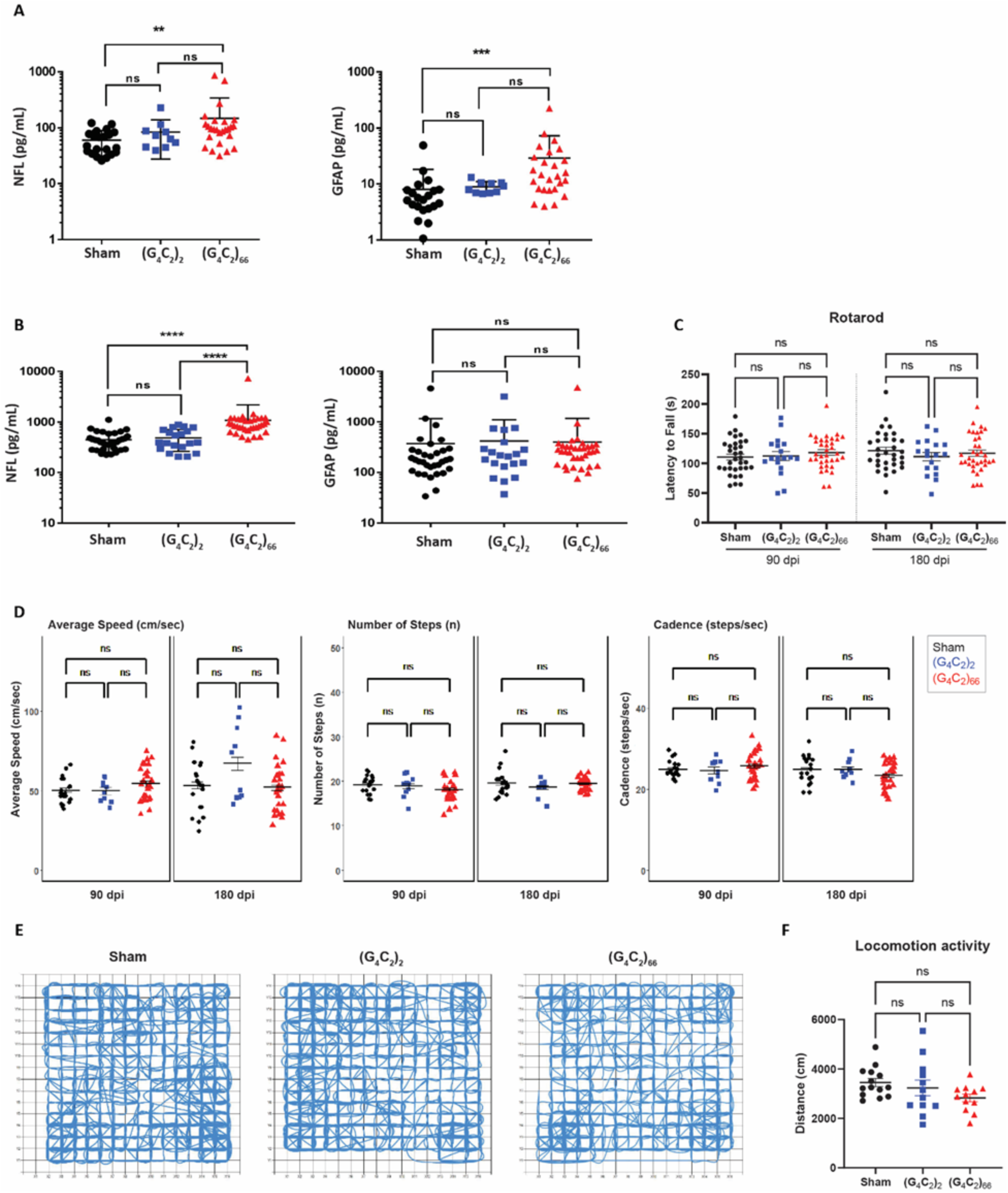
(G_4_C_2_)_66_ mice do not have behavioral deficits. Analysis of NfL and GFAP levels in plasma **(A)** (sham n= 21, (G_4_C_2_)_2_ n= 10, (G_4_C_2_)_66_ n= 26). One way ANOVA followed by Dunnett’s test (Dunnett, 1955). (** p < 0.01, *** p < 0.001 vs sham), and CSF **(B)** (sham n= 33, (G_4_C_2_)_2_ n= 20, (G_4_C_2_)_66_ n= 35). One way ANOVA followed by Dunnett’s test. (**** p < 0.0001 vs sham), error bars = SEM **(C)** Rotarod behavioral test at 90 dpi and 180 dpi (sham n=33, (G_4_C_2_)_2_ n=18, (G_4_C_2_)_66_ n=35). A repeat-measure two-way ANOVA analysis (Time: p=0.7403; Time x Group: p=0.2446; Group: 0.7403) with Sidak’s multiple comparison test was performed on the dataset. 90 dpi: sham vs. (G_4_C_2_)_2_ p=0.9995, sham vs. (G_4_C_2_)_66_ p=0.7333, (G_4_C_2_)_2_ vs. (G_4_C_2_)_66_ p=0.8722; 180 dpi: sham vs. (G_4_C_2_)_2_ p=0.5913, sham vs. (G_4_C_2_)_66_ p=0.9142, (G_4_C_2_)_2_ vs. (G_4_C_2_)_66_ p=0.8827, error bars = SEM. **(D)** Catwalk behavioral test at 90 dpi and 180 dpi (sham n=20, (G_4_C_2_)_2_ n=10, (G_4_C_2_)_66_ n=27), including: average speed (cm/sec), number of steps (n), and cadence (steps/sec). Linear mixed-effects model was performed with least-squares means compared among groups and multiplicity controlled with Tukey’s multiple comparisons test. Average speed: sham vs. (G_4_C_2_)_2_ p=1.000 (90 dpi), p=0.9579 (180 dpi); sham vs. (G_4_C_2_)_66_ p=0.9975 (90 dpi), p=1.000 (180 dpi); (G_4_C_2_)_2_ vs. (G_4_C_2_)_66_ p=1.000 (90 dpi), p=0.8401 (180 dpi). Number of steps: sham vs. (G_4_C_2_)_2_ p=1.000 (90 dpi), p=1.000 (180 dpi); sham vs. (G_4_C_2_)_66_ p=0.9957 (90 dpi), p=1.000 (180 dpi); (G_4_C_2_)_2_ vs. (G_4_C_2_)_66_ p=1.000 (90 dpi), p=1.000 (180 dpi). Cadence: sham vs. (G_4_C_2_)_2_ p=1.000 (90 dpi), p=1.000 (180 dpi); sham vs. (G_4_C_2_)_66_ p=1.000 (90 dpi), p=0.9997 (180 dpi); (G_4_C_2_)_2_ vs. (G_4_C_2_)_66_ p=1.000 (90 dpi), p=1.000 (180 dpi), error bars = SEM. **(E)** Representative paths of sham, (G_4_C_2_)_2_, and (G_4_C_2_)_66_ injected mice in open-field behavioral test at 180 dpi. **(F)** Analysis of locomotion activity in the open-field test at 180 dpi (sham n=14, (G_4_C_2_)_2_ n=12, (G_4_C_2_)_66_ n=12). One-way ANOVA analysis of locomotion activity was performed (p=0.1406) followed by Tukey’s multiple comparison test. sham vs (G_4_C_2_)_2_ p=0.7597, sham vs (G_4_C_2_)_66_ p=0.1219 and (G_4_C_2_)_2_ vs (G_4_C_2_)_66_ p=0.4241, error bars = SEM.

### C9ORF72 (G4C2)66 mice do not have notable behavioral deficits

We evaluated the behavioral effects of the (G_4_C_2_)_66_ repeat expansion with various behavioral tests including rotarod, grip strength, catwalk, and open-field assessment at 6-month of age. The study was powered based on rotarod as the primary end point, which indicated that the initial target sample size of n=20 per group would provide 80% power to find significant differences as small as 23.3 seconds between groups on rotarod (see Randomization, Blinding, and Power section for details). There was no significant effect on bodyweight in (G_4_C_2_)_66_ males from 90 dpi to 180 dpi compared to controls, while a slight decrease was notable in (G_4_C_2_)_66_ females starting 146 dpi (**Supplemental Figure 5A**). We assessed rotarod performance at 90 dpi and 180 dpi and found no significant difference between the experimental groups (**Figure 7C**). After accounting for attrition among cohorts during the study, a post-hoc power analysis revealed that the final sample sizes (n=32 Sham, n=18 (G_4_C_2_)_2_ and n=35 (G_4_C_2_)_66_) were sufficient to provide at least 80% power to detect a difference as small as 25 seconds in latency to fall on rotarod when comparing Sham or (G_4_C_2_)_66_ to (G_4_C_2_)_2_, and a difference as small as 20 seconds when comparing Sham to (G_4_C_2_)_66_. Given that the observed differences among the groups on this study were much smaller than 20 seconds, we can conclude that the lack of statistical significance among groups in the rotarod assay is not due to an insufficient sample size. We evaluated forelimb and hindlimb grip strength in females and males at 90 dpi and no significant difference was detected between groups (**Supplemental Figure 5B**). Gait was assessed in the animals using Catwalk analysis, where no significant change to average speed, number of steps, or cadence was evident between the experimental groups (**Figure 7D**). Finally, we used the open-field assay to evaluate general locomotor activity, exploration, and anxiety-like behavior in a novel environment at 180 dpi (**Figure 7E-F; Supplemental Figure 5C**). There was no obvious or significant difference between the groups. Collectively, we did not observe any behavioral changes in the (G_4_C_2_)_66_ animals unlike the previously reported study [10].

## Discussion

ALS is a complex neurodegenerative disease that demonstrates regional specificity at onset and during progression along with highly consistent molecular pathologies [62]. Although sALS accounts for a significant portion (>85%) of all cases, fALS has proven relatively more straightforward to study because of patient-derived induced pluripotent stem cells (which harbor the endogenous disease-causing mutations) and animal models that have been genetically modified to express disease-causing mutations in evolutionarily conserved genes [50, 63]. However, it is important to note that introduction of a mutation, or overexpression of human disease-associated proteins in non-human animals may not necessarily recapitulate all the physiological or pathological hallmarks of human disease. Here, we have evaluated histological, behavioral, and biochemical endpoints to determine which endophenotypes of human ALS/FTD are recapitulated in a previously published AAV2/9 *C9ORF72* (G_4_C_2_)_66_ overexpression model [10] using a robust study design.

Firstly, this injection-based model leads to an extensive transduction of neurons throughout the brain and rostral segments of the spinal cord. We were able to detect the virally expressed repeat RNA in most brain regions and in cervical spinal cord but limited expression in lumbar spinal cord (refer to **Supplemental Figure 1B**). We interrogated the expression of polyGP, one of the most abundant DPRs translated from repeat RNA, via MSD assay and immunohistochemistry. The polyGP MSD assay provided a sensitive but bulk assessment of brain and thoracic spinal cord samples, where polyGP burden was significantly elevated in the (G_4_C_2_)_66_ animals. However, the burden of polyGP in thoracic spinal cord samples was approximately ten times less than that detected in the brain. Concordantly, there was notably lower poly-GP detected by immunohistochemistry in cervical spinal cord relative to the brain and even lower in the lumbar spinal cord, further corroborating a rostral-to-caudal decrease in viral expression and subsequent DPR expression. The other two DPRs, polyGA and polyGR, were detectable in the brain, but only minimally evident in the cervical spinal cord. Some polyGA and polyGR accumulations may be present in the lumbar spinal cord, though we were unable to faithfully detect the inclusions due to high background signal in control sections (data not shown). The decreasing trend in DPR abundance moving caudally throughout the spinal cord matches our observations that the repeat RNA is barely detectable in the lumbar spinal cord, though it is clearly visualized in the cervical region.

This observation, perhaps unsurprisingly, led us to question whether the (G_4_C_2_)_66_ mice exhibit motor deficits, given the low abundance of repeat RNA and DPRs in the spinal cord. It has previously been reported that mice injected with the *C9ORF72* (G_4_C_2_)_66_ viral construct fail to improve in daily rotarod tests relative to controls [10]. The authors did not observe motor deficits on the initial day of testing, yet a significant difference between (G_4_C_2_)_66_ and control mice becomes evident after more than 2 days of consecutive rotarod tests. However, it is important to note that there was no baseline assessment prior to these tests. In this study, we assessed latency to fall during a single day of rotarod testing at 90 dpi and 180 dpi with a well-powered sample size. There was no significant difference in the rotarod performance of (G_4_C_2_)_66_ mice relative to sham or (G_4_C_2_)_2_ controls. Moreover, we assessed various other metrics of motor function including grip strength and gait analysis and found no detectable difference in (G_4_C_2_)_66_ mice compared to control groups. Given the lower burden of repeat RNA and DPRs in the spinal cord, we do not believe these mice demonstrate any primary deficits in motor function. However, it is possible that the high burden of repeat and DPRs in the motor cortex leads to a reduction in learned motor ability, which would become evident following multiple days of rotarod testing.

Additionally, it would be intriguing to assess this model for memory deficits—as the significant DPR burden in the brain but limited expression in the spinal cord may have a greater impact on neurons involved in memory formation and retention.

Although there was no overt cellular loss observed in the CNS, there were hallmarks of cellular injury detected the(G_4_C_2_)_66_ animals. The increase in GFAP certainly reflects an astroglial response to cellular injury. Furthermore, the elevated NfL also is evidence for some neuronal injury as well. Nevertheless, the extent of this CNS injury was not sufficient to impair the various overall motor function or other behavioral measures, nor was it sufficient to be linked to any loss of TDP-43 function.

While loss of motor function is a consistent clinical characteristic of patients with ALS, there are many cellular and molecular signatures that allow clearer diagnosis postmortem. For example, nuclear clearance and loss of nuclear function of TDP-43, a nuclear DNA- and RNA-binding protein, is one of the most prominent pathological events preceding neuronal death [66, 70]. This pathology is present in the vast majority of ALS cases (>97%) and is often co-observed with p62-positive granules [35]. We were unable to observe any significant nuclear clearance of TDP-43 in neurons of the motor cortex, despite the presence of p62-postive granules in this region. Since loss of TDP-43 nuclear function might occur prior to observed nuclear clearance by immunofluorescent assessment, we interrogated TDP-43 nuclear function by quantifying the abundance of multiple mRNA targets regulated by TDP-43. None of these targets, identified in previously published RNA-sequencing studies [86], demonstrated any significant alterations due to the presence of the *C9ORF72* (G_4_C_2_)_66_ construct. This was true in both the thoracic spinal cord, where there is low viral burden, and the brain, which bears significant viral burden. Additionally, we observed no significant presence of phosphorylated TDP-43, a proposed pathogenic form, despite attempts with numerous commercial and custom pTDP-43 (Ser409/410) antibodies (data not shown) and no mislocalization of TDP-43 in cells positive for DPRs. Taken together, these findings provide evidence that, at least in the C57BL/6J mouse brain, overexpression of DPRs and repeat RNA in neurons does not lead to TDP-43 pathology or loss of function in 6-month-old (180 dpi) animals. Perhaps broader introduction of the virus to all cells within the central nervous system may lead to aberrant TDP-43 pathology in neurons via non-cell autonomous effects, as this would more closely match the presence of the expanded repeat in human patients. Alternatively, this may be characteristic of interspecies differences.

There are more than ninety mouse models that have been generated to aid in the study of ALS [15]. Although there is significant redundancy when it comes to the genes being studied (for example, there are over thirty models to study TDP-43 alone), each model demonstrates unique strengths and weaknesses. While the *C9ORF72* (G_4_C_2_)_66_ viral model we use in this study exhibits neither TDP-43 pathology nor primary motor deficits, the abundant RNA and DPR pathology makes it a useful tool to understand these pathological species’ biogenesis, degradation, molecular associations, and traffic within and between cells. Use of this viral model would greatly complement studies that recapitulate more prominent aspects of ALS but fail to produce DPRs or repeat RNA at high enough levels to reliably investigate their pathology. Since ALS is such a heterogeneous disease in terms of genetic cause, onset, and progression, perhaps it is prudent to use multiple distinct models in each investigation. Though this will prove to be more costly and time consuming, it may lead to more reliable findings and results that ultimately represent human disease.

## Supplemental Figures

**Supplemental Figure 1.**
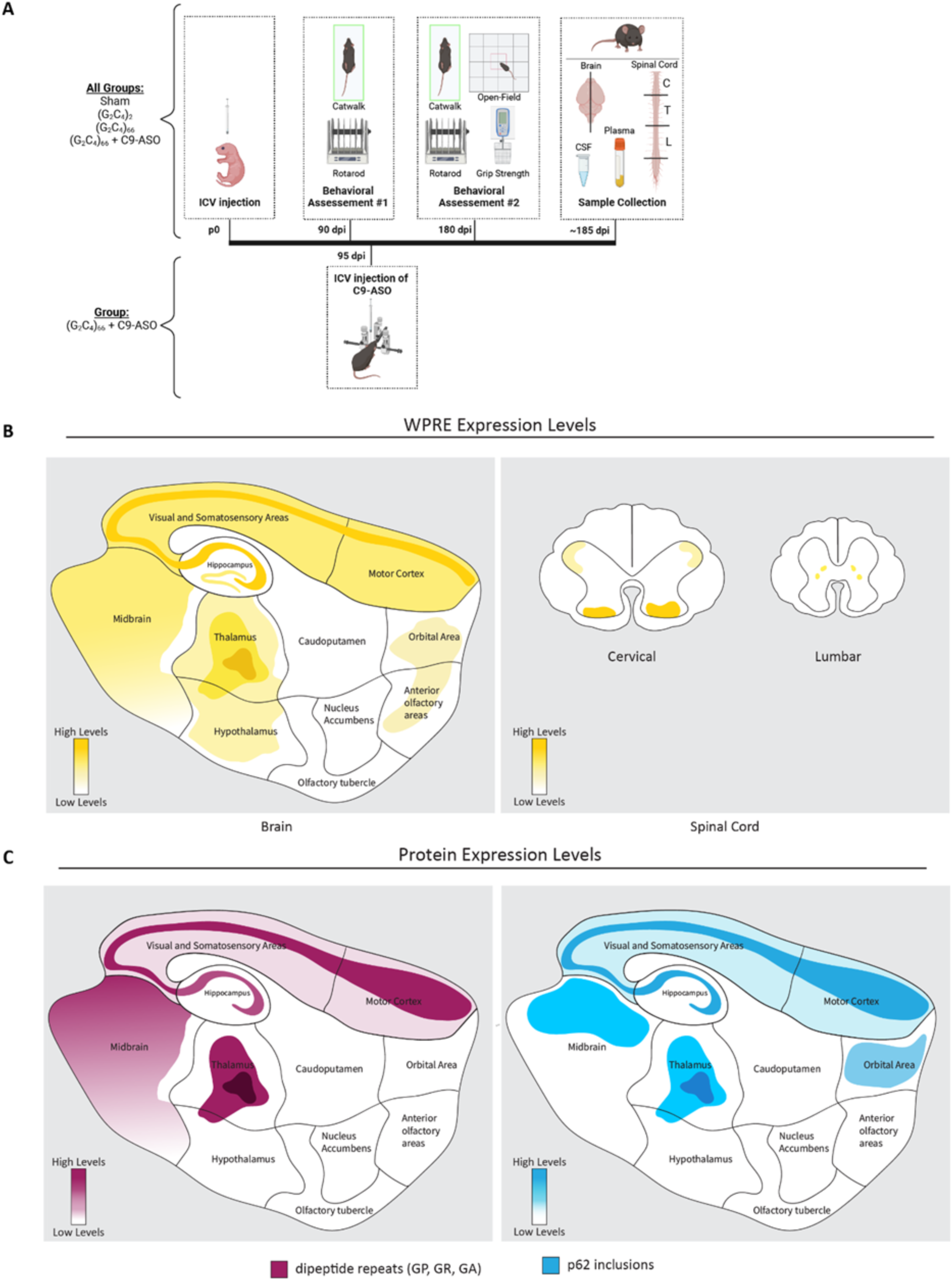
**(A)** Schematic overview of the study design. Cartoon representation of spatial viral RNA expression **(B)** and protein expression levels of dipeptide repeat expression (purple) and p62 inclusions (blue) in **(C)**. Both DPR expression and p62 inclusions are localized to the areas where viral (G_4_C_2_)_66_ RNA is expressed.

**Supplemental Figure 2.**
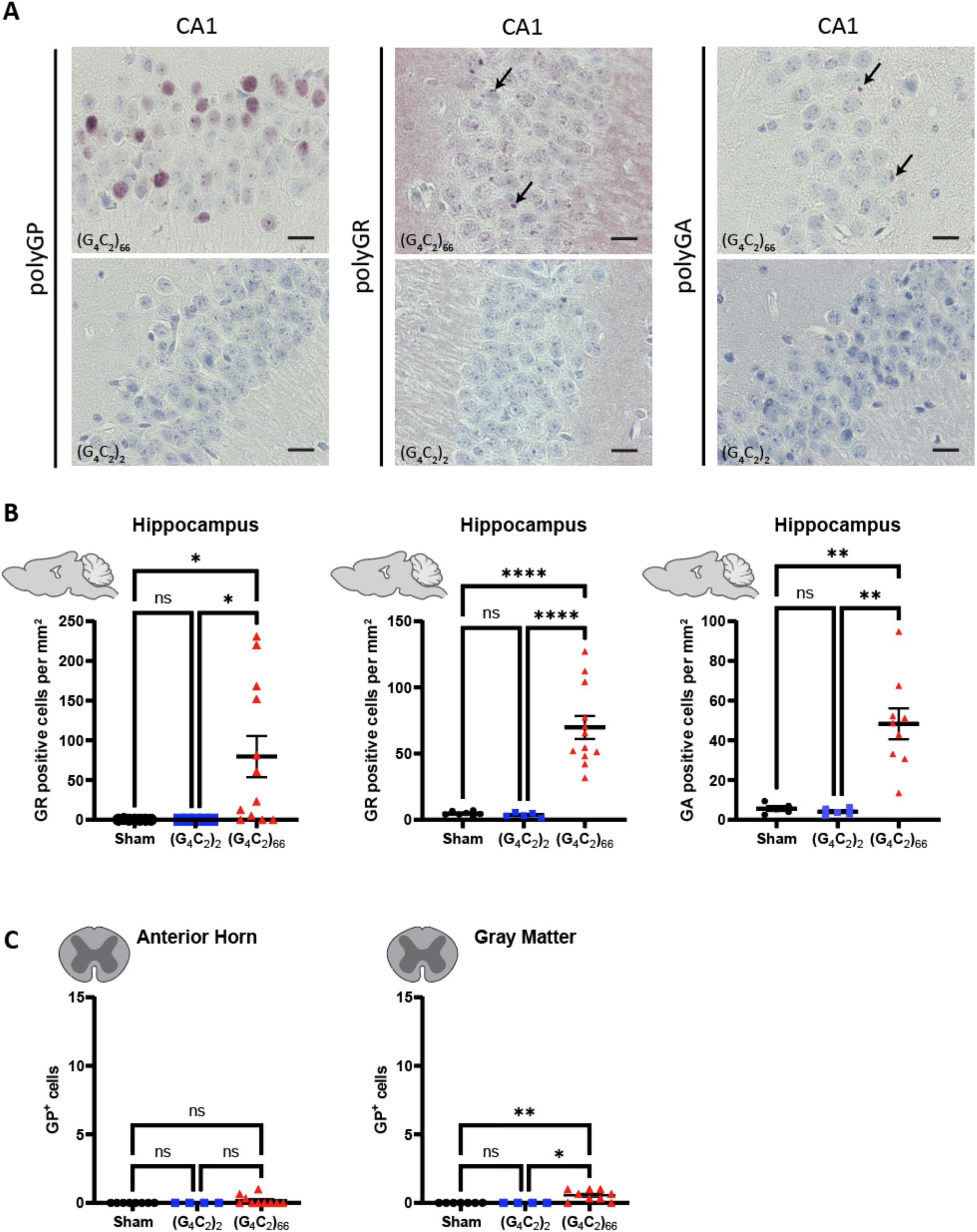
(G_4_C_2_)_66_ mice have DPR expression in the hippocampus. **(A)** Representative images of CA1 region from brain sections stained against polyGP, polyGR, and polyGA. Scale bars 20 µm. **(B)** Quantification of sections stained against polyGP, polyGR, and polyGA (sham n=14, (G_4_C_2_)_2_ n=12, (G_4_C_2_)_66_ n=12). One-way Welch ANOVA analysis of polyGP, polyGR, and polyGA positive cell number was performed (p=0.0021 for polyGP, p<0.0001 for polyGR, p=0.0001 for polyGA) followed by Tukey’s multiple comparison test. polyGP: sham vs (G_4_C_2_)_2_ p=0.9993, sham vs (G_4_C_2_)_66_ p=0.0303 and (G_4_C_2_)_2_ vs (G_4_C_2_)_66_ p=0.0304, polyGR: sham vs (G_4_C_2_)_2_ p=0.6542, sham vs (G_4_C_2_)_66_ p<0.0001 and (G_4_C_2_)_2_ vs (G_4_C_2_)_66_ p<0.0001, polyGA: sham vs (G_4_C_2_)_2_ p=0.6636, sham vs (G_4_C_2_)_66_ p=0.0018 and (G_4_C_2_)_2_ vs (G_4_C_2_)_66_ p=0.0013, error bars = SEM. **(C)** Quantification of lumbar spinal cord sections stained against polyGP (sham n=7, (G_4_C_2_)_2_ n=4, (G_4_C_2_)_66_ n=9). A Kruskal-Wallis test of the lumbar spinal cord gray matter (p=0.0015) and anterior horn (p=0.1870) dataset was performed, followed by Dunn’s multiple comparisons test (gray matter: sham vs. (G_4_C_2_)_2_ p>0.9999, sham vs. (G_4_C_2_)_66_ p=0.0064, (G_4_C_2_)_2_ vs. (G_4_C_2_)_66_ p=0.0301; anterior horn: sham vs. (G_4_C_2_)_2_ p>0.9999, sham vs. (G_4_C_2_)_66_ p=0.2180, (G_4_C_2_)_2_ vs. (G_4_C_2_)_66_ p=0.4504), error bars = SEM.

**Supplemental Figure 3.**
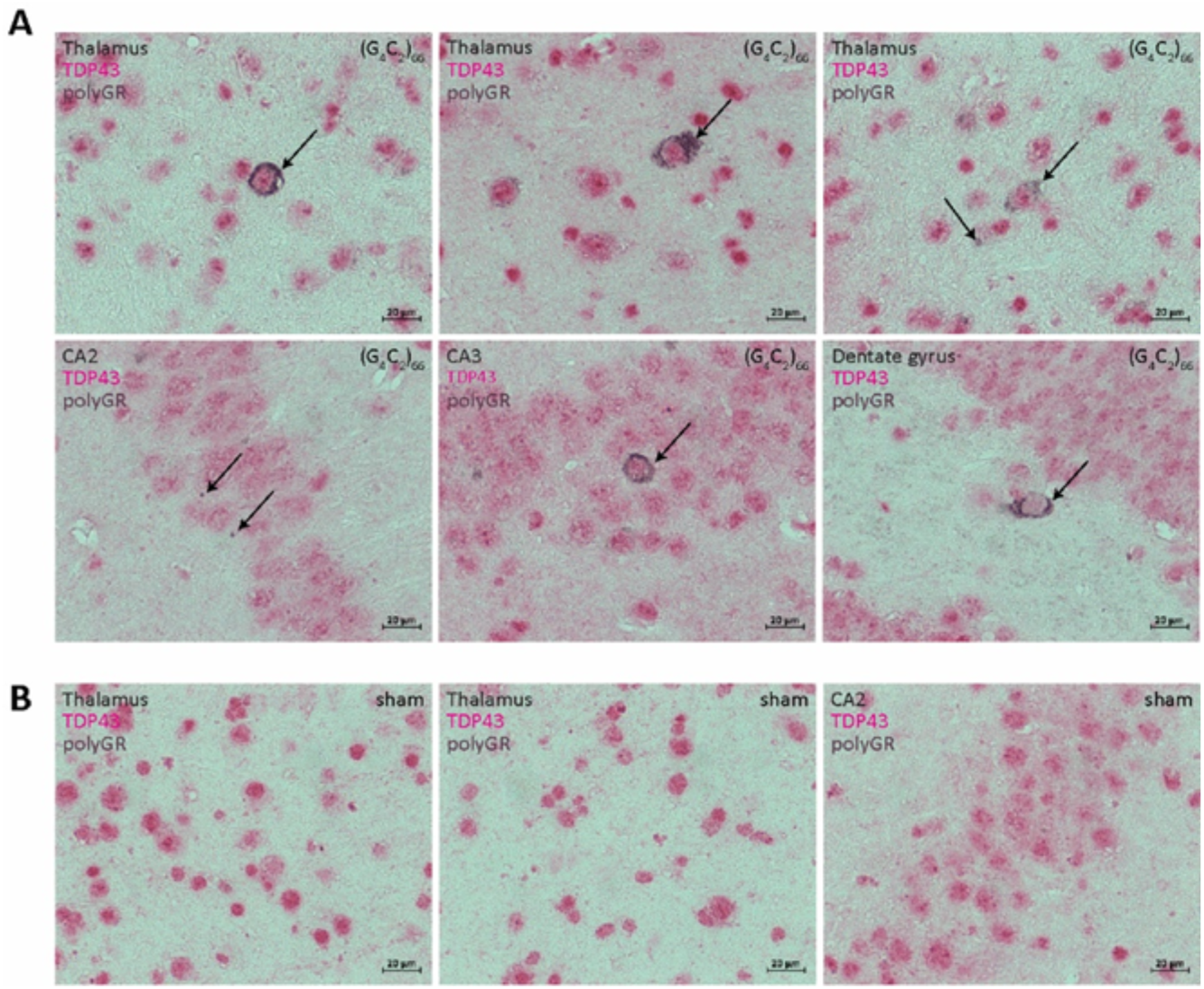
TDP-43 does not mislocalize in polyGR positive cells. **(A)** Representative images of Thalamus, CA1, CA3, and Dentate Gyrus brain regions in (G_4_C_2_)_66_ mice co-labled with TDP-43 (pink) and polyGR (brown). Scale bars 20 µm. No mislocalization of TDP-43 into the cytoplasm is observed in cells positive for TDP-43 and polyGR (denoted by black arrows). **(B)** Representative images of Thalamus and CA2 brain regions in sham mice co-labled with TDP-43 (pink) and polyGR (brown). No polyGR positive cells are present in sham tissue. Scale bars 20 µm.

**Supplemental Figure 4.**
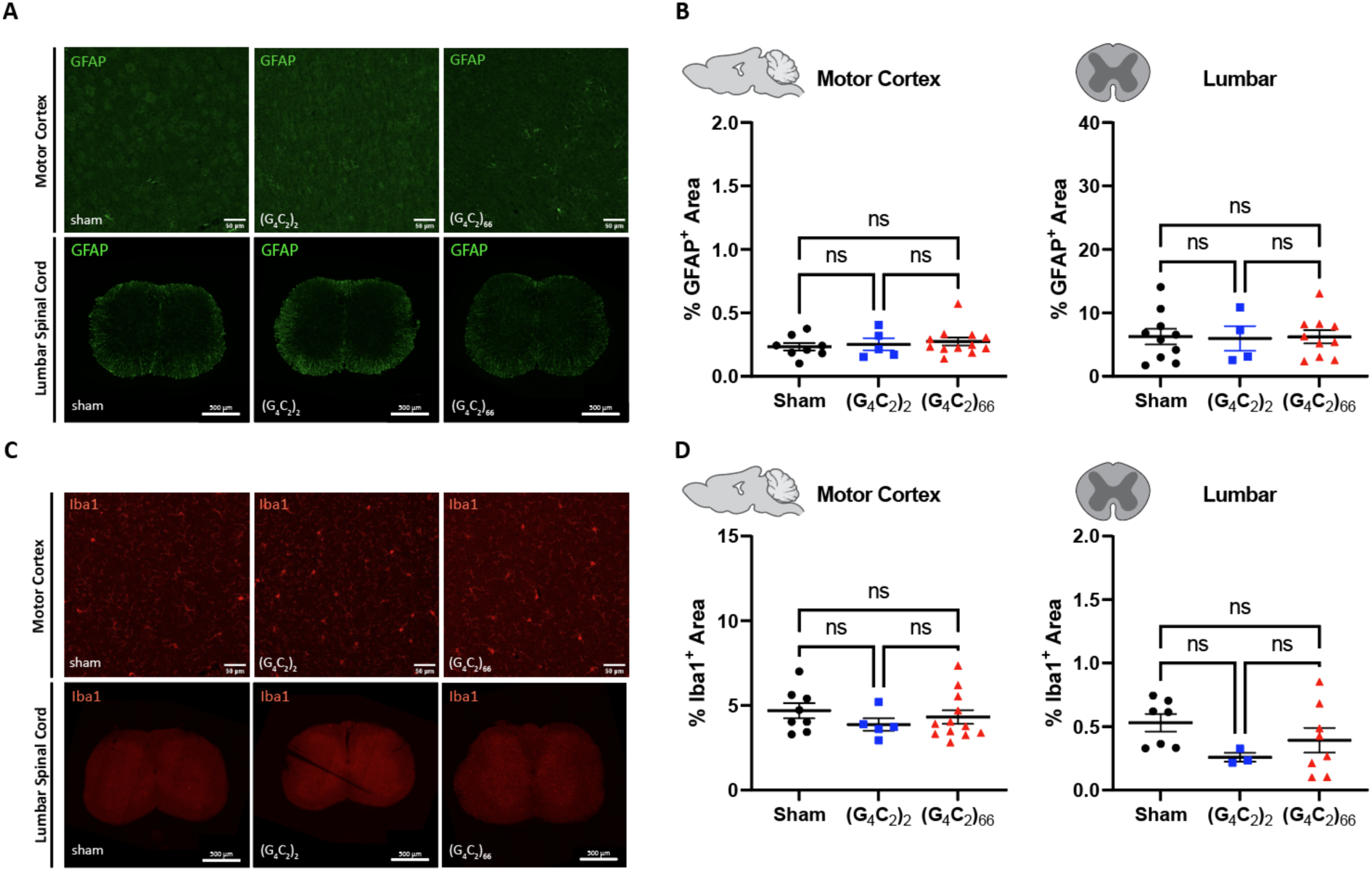
Analysis of gliosis in the (G_4_C_2_)_66_ mouse model by immunohistochemistry. **(A)** Representative images of immunohistochemistry analysis of GFAP expression within the motor cortex and lumbar spinal cord. Motor cortex scale bars 50 µm and lumbar spinal cord scale bars 200 µm. **(B)** Quantification of % GFAP positive area within cortex and cervical spinal cord (sham n=8 for motor cortex and 10 for spinal cord, (G_4_C_2_)_2_ n=5 for, motor cortex and 4 for spinal cord, (G_4_C_2_)_66_ n=12 for motor cortex and 10 for spinal cord). One-way ANOVA analysis of the motor cortex dataset was performed (p=0.6690), followed by Tukey’s multiple comparisons test: sham vs. (G_4_C_2_)_2_ p=0.9433, sham vs. (G_4_C_2_)_66_ p=0.6479, (G_4_C_2_)_2_ vs. (G_4_C_2_)_66_ p=0.9077. One-way ANOVA analysis of the lumbar spinal cord dataset was performed (p=0.9891), followed by Tukey’s multiple comparisons test: sham vs. (G_4_C_2_)_2_ p=0.9888, sham vs. (G_4_C_2_)_66_ p=0.9998, (G_4_C_2_)_2_ vs. (G_4_C_2_)_66_ p=0.9909, error bars = SEM. **(C)** Representative images of immunohistochemistry analysis of Iba1 expression within motor cortex and lumbar spinal cord. Cortex scale bars 50 µm and lumbar spinal cord scale bars 200 µm. **(D)** Quantification of % Iba1 positive area within motor cortex and lumbar spinal cord (sham n=8 for motor cortex and 7 for spinal cord, (G_4_C_2_)_2_ n=5 for motor cortex and 3 for spinal cord, (G_4_C_2_)_66_ n=12 for motor cortex and 8 for spinal cord). One-way ANOVA analysis of the motor cortex dataset was performed (p=0.5259), followed by Tukey’s multiple comparisons test: sham vs. (G_4_C_2_)_2_ p=0.4978, sham vs. (G_4_C_2_)_66_ p=0.7929, (G_4_C_2_)_2_ vs. (G_4_C_2_)_66_ p=0.7852. One-way ANOVA analysis of the lumbar spinal cord dataset was performed (p=0.2099), followed by Tukey’s multiple comparisons test: sham vs. (G_4_C_2_)_2_ p=0.2085, sham vs. (G_4_C_2_)_66_ p=0.4674, (G_4_C_2_)_2_ vs. (G_4_C_2_)_66_ p=0.6513, error bars = SEM.

**Supplemental Figure 5.**
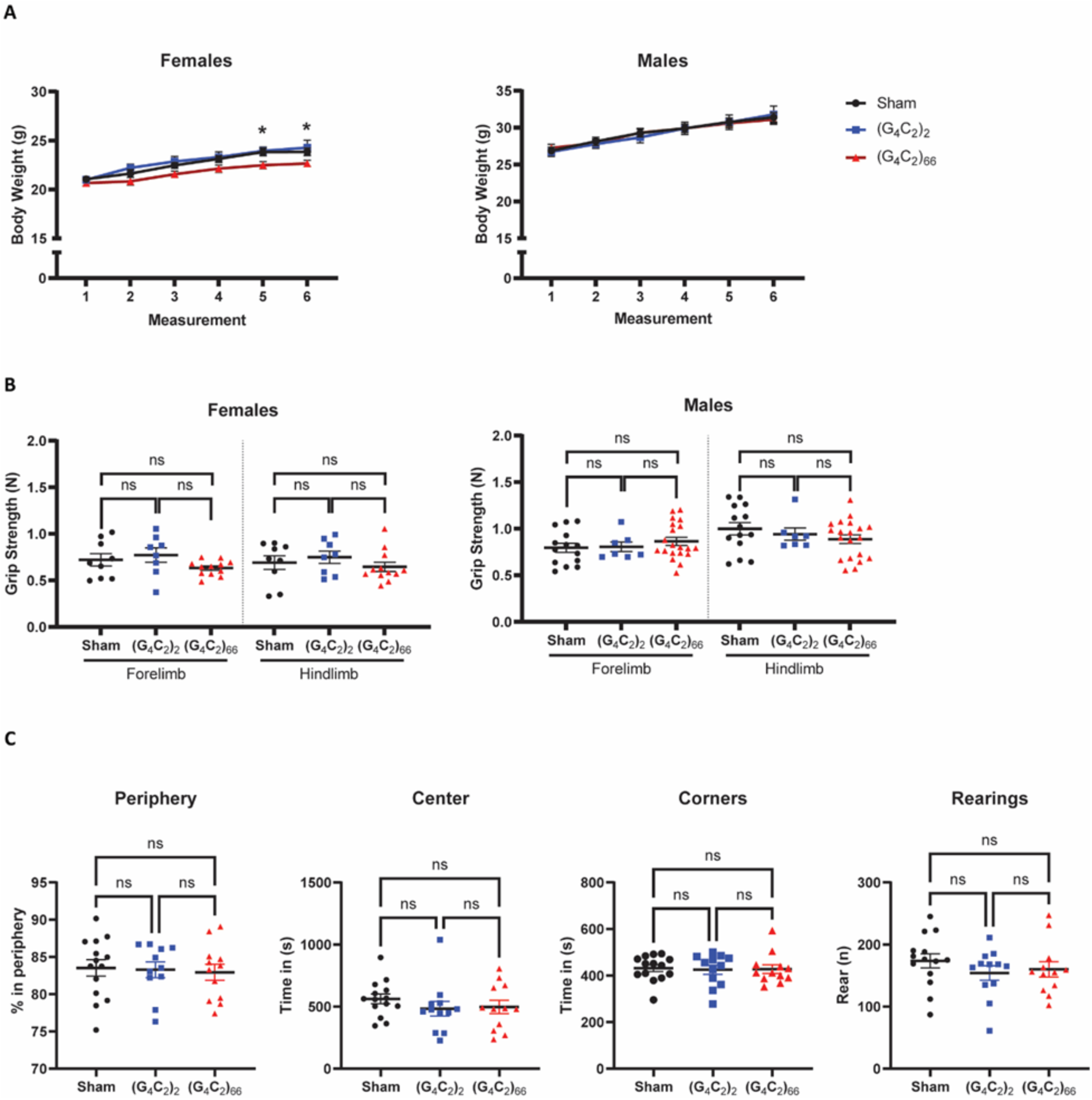
(G_4_C_2_)_66_ mice do not have behavioral deficits. **(A)** Bodyweight tracking of mice (females: sham n=18, (G_4_C_2_)_2_ n=8, (G_4_C_2_)_66_ n=15, males: sham n=14, (G_4_C_2_)_2_ n=10, (G_4_C_2_)_66_ n=20). A two-way repeated-measures ANOVA analysis of the female dataset was performed (p=0.0573), followed by Dunnett’s multiple comparisons test where a significant difference was observed at the fifth (sham vs. (G_4_C_2_)_66_ p=0.0276) and sixth sham vs. (G_4_C_2_)_66_ p=0.0479 measurement. A two-way repeated-measures ANOVA analysis of the male dataset was performed (p=0.9917), followed by Dunnett’s multiple comparisons test but no significant difference was found between the groups for any of the bodyweight measurements, error bars = SEM. **(B)** Grip strength behavioral analysis at 90 dpi in female (sham n=9, (G_4_C_2_)_2_ n=8, (G_4_C_2_)_66_ n=12) and male mice (sham n=14, (G_4_C_2_)_2_ n=7, (G_4_C_2_)_66_ n=20). One-way ANOVA analysis of the female dataset was performed (forelimb: p=0.1871, hindlimb: p=0.5037), followed by Tukey’s multiple comparisons test: sham vs. (G_4_C_2_)_2_ (forelimb: p=0.8087, hindlimb: p=0.8097), sham vs. (G_4_C_2_)_66_ (forelimb: p=0.4616, hindlimb: p=0.8514), (G_4_C_2_)_2_ vs. (G_4_C_2_)_66_ (forelimb: p=0.1808, hindlimb: p=0.4722). One-way ANOVA analysis of the male dataset was performed (forelimb: p=0.5226, hindlimb: p=0.3495), followed by Tukey’s multiple comparisons test: sham vs. (G_4_C_2_)_2_ (forelimb: p=0.9918, hindlimb: p=0.8354), sham vs. (G_4_C_2_)_66_ (forelimb: p=0.5329, hindlimb: p=0.3175), (G_4_C_2_)_2_ vs. (G_4_C_2_)_66_ (forelimb: p=0.7492, hindlimb: p=0.8414), error bars = SEM. **(C)** Time spent in periphery, center, corners and number of rearing analyses in open-field test at 180 dpi (n=14 sham, n=12 (G_4_C_2_)_2_, n=12 (G_4_C_2_)_66_). One-way ANOVA analyses of time spent in periphery, in center, in corners, and the number of rearing were performed (p=0.9207 for time spent in periphery, p=0.4894 for time spent in center, p=0.9647 for time spent in corners, and p=0.4693 for number of rearings) followed by Tukey’s multiple comparison test. Periphery: sham vs (G_4_C_2_)_2_ p=0.9840, sham vs (G_4_C_2_)_66_ p=0.9130 and (G_4_C_2_)_2_ vs (G_4_C_2_)_66_ p=0.9741, Center: sham vs (G_4_C_2_)_2_ p=0.5099, sham vs (G_4_C_2_)_66_ p=0.6300 and (G_4_C_2_)_2_ vs (G_4_C_2_)_66_ p=0.9807, Corners: sham vs (G_4_C_2_)_2_ p=0.9620, sham vs (G_4_C_2_)_66_ p=0.9864 and (G_4_C_2_)_2_ vs (G_4_C_2_)_66_ p=0.9941, Rearings: sham vs (G_4_C_2_)_2_ p=0.4598, sham vs (G_4_C_2_)_66_ p=0.6833 and (G_4_C_2_)_2_ vs (G_4_C_2_)_66_ p=0.9339, error bars = SEM.

## Contributions

E.G.T., O.S., C.C., T.P., G.N., J.D.R. designed the experiments. C.C. provided blinding, randomization, and power analysis for this study. E.G.T., O.S., S.C.A., E.R.K., T.P., B.G.V., W.Z., L.X., conducted all in-life experimentation. E.G.T., O.S., S.C.A., B.L.Z. performed all molecular, biochemical, and histological experimentation. A.Z. conducted the analysis of NfL and GFAP concentrations in plasma and CSF. S.C.A. performed statistical analysis of open-field and p62 and DPR histology. C.C. performed statistical analysis of catwalk data. E.G.T. performed all other statistical analysis in this study. E.G.T., O.S., C.C., S.C.A., B.L.Z., G.N. wrote the paper. All authors reviewed and edited this manuscript.

## Acknowledgements

This work was funded in large part by a Research Collaboration Agreement between JHU and GSK. Additional funding came from the Robert Packard Center for ALS Research, NIH-NINDS (R01 5R35NS132179-02, J.D.R and F32NS120940, E.G.T.), and the ALS Association Milton Safenowitz Postdoctoral Fellowship (E.G.T. and S.C.A.).

